# A newly arisen indel governs a leaf shape polymorphism in the Ivy Leaf Morning Glory (*Ipomoea hederacea*)

**DOI:** 10.64898/2026.06.04.730136

**Authors:** Amanda L. Peake, Emily Glasgow, Claire Abbasi, Yunchen Gong, Lauren Whitt, Melissa Williams, Jane Grimwood, Alex Harkess, John R. Stinchcombe

## Abstract

Leaf shape varies widely across plant taxa and has repeatedly been shown to affect ecophysiology, interspecific interactions, and fitness. We used population genomics, genome wide association studies (GWAS), and comparative genomics to determine the genetic basis and evolutionary history of an uncharacterized Mendelian leaf shape polymorphism in *Ipomoea hederacea*. To do so, we assembled a reference genome and generated whole genome sequencing for 123 individuals from 55 populations. We identified a 117 kb indel that perfectly co-segregates with leaf shape by conducting a GWAS and assessing differences in coverage. Syntenic orthologs for genes on the indel were present in five other *Ipomoea* species with various leaf shapes, indicating the indel is a newly arisen deletion in *I. hederacea* despite similar leaf shape phenotypes in the other species.

More broadly, these results illustrate how a range of leaf shape phenotypes can be produced by distinct genetic mechanisms even in closely related species. Although none of the genes on the indel itself are known leaf shape candidates, there are multiple leaf shape candidate genes in close proximity that are involved in the auxin biosynthesis pathway. Additionally, the genes within the indel have gene functions that could affect other potentially ecologically relevant traits that have previously been shown to be associated with leaf shape in *I. hederacea*. Therefore, the pleiotropic effects of the indel polymorphism can have important implications for understanding the ecological mechanisms influencing a well-documented leaf shape latitudinal cline in *I. hederacea*.

**Significance Statement:** *Ipomoea hederacea* has been used to investigate the ecological and evolutionary effects of leaf shape because of a well-documented latitudinal leaf shape cline governed by an uncharacterized Mendelian polymorphism in this species. We identified a 117 kb indel that perfectly co-segregates with leaf shape in *I. hederacea*. The indel appears to be a distinct genetic mechanism than those governing similar leaf shapes in other closely related *Ipomoea* species. Additionally, possible pleiotropic effects of the indel on other traits has important implications for understanding the ecology and evolution of leaf shape variation in *I. hederacea*. Therefore, the results and the new genomic resources that we developed will facilitate future work understanding the genetic, developmental, and ecological mechanisms governing leaf shape variation.

## Introduction

Why are leaves the shape they are? Evolution has given rise to a tremendous amount of leaf morphology where even slight differences in leaf shape can have major functional, ecological, and evolutionary ramifications (1). Differences in leaf shape are not only observed at the species level, but distinct leaf shapes have often been documented between populations and individuals of the same species (2–5). Although leaf shape can sometimes be hard to quantify, there are frequently measurable differences in length-to-width ratios (6), number of teeth or serrations (7), and number or degree of lobing (8–10). Considering that leaves perform many important functions, characterizing the genetic basis of leaf shape can directly inform how evolutionary forces and ecological factors together maintain phenotypic variation (1). For example, if a leaf shape locus has pleiotropic effects, adaptive evolution in leaf shape may impact phenotypic evolution of other ecologically relevant traits, and vice versa. Similarly, the repeated evolution of a phenotype by the same genetic mechanisms indicates relatively few genetic routes for evolution and is thought to be an indicator of pleiotropy (11–13), whereas independent genetic mechanisms suggest little evolutionary constraint, and multiple genetic and developmental means of producing phenotypes. Here, we describe the genetic basis of a Mendelian leaf shape polymorphism in *Ipomoea hederacea* and surprisingly, we show that a 117 kb deletion governing the polymorphism is likely newly arisen in the species despite similar leaf shape phenotypes in other related *Ipomoea* species.

The *Ipomoea* genus displays remarkable phenotypic variation in flower color, growth forms, and leaf shapes, amongst other ecologically important traits (14). Combined with the ongoing development of *Ipomoea* genomic resources, the genus provides an exemplary study system for ecological and comparative genomic research. Leaf shape varies widely within the *Ipomoea* genus across multiple taxonomic levels. At the species level, leaf shape ranges from some species that almost exclusively have entire-shaped leaves, such as *I. purpurea*, to other species with deep pinnately-lobed leaves, such as *I. quamoclit* (15). Leaf shape variation also occurs within and between populations in species like *I. hederacea* (4, 8) and *I. batatas* (16, 17). The genetic basis of leaf shape also differs between congeneric species, for example, lobing in *I. hederacea* is governed by a single Mendelian polymorphism (8, 18) whereas multiple candidate loci and GxE have been shown to impact leaf shape in *I. batatas* (16, 17). Additionally, cultivated ornamental *I. nil* lines have striking variation in leaf shape governed by genes mapped to different linkage groups (19–22). By identifying the genetic mechanisms governing leaf shape differences within a species we can then determine if the genetic mechanism is shared amongst other species or if it is a newly derived evolutionary innovation. A better understanding of how mutational effect-sizes and pleiotropy could be influencing leaf shape evolution requires knowing whether similar leaf shapes have evolved repeatedly by mutations within different genes or by multiple independent mutations within the same gene (12).

The *Ipomoea* genus provides an opportunity to leverage both intra- and inter-specific diversity to understand how the genetic basis and evolution of leaf shape influence each other.

Across the Eastern United States, *I. hederacea* has a latitudinal leaf shape cline governed by a single, uncharacterized Mendelian polymorphism (3, 4, 8). Homozygotes for alternative alleles have either lobed or entire-shaped leaves (Figure 1A) and heterozygotes– which are rare in natural populations– have an intermediate phenotype (8, 18). Despite the simple genetic basis, the leaf shape locus has not yet been identified due to the lack of a genome assembly and high resolution genotyping for the species. Across the North American species range, lobed individuals are more prominent in northern populations and entire-shaped individuals are more common in southern populations (3, 4, 8). Herbarium and contemporary collections suggest that multiple phenotypic clines are being maintained (4, 8, 23, 24) despite the overall lack of geographic population genetic structure in *I. hederacea* (4, 25), suggesting that the latitudinal clines reflect adaptive phenotypic variation. Moreover, the leaf shape locus is an outlier when compared to putatively neutral loci (4, 25). The alternative leaf shape genotypes in *I. hederacea* have been shown to differ in night-time thermoregulation (26), stomatal conductance in the field on sunny days (27), herbivory preferences (28), pathogen resistance (3), and leaf microbiome assembly (29). Previous population genetic studies in *I. hederacea*, however, have been limited to genetic datasets using anonymous AFLP markers (4) or Sanger sequencing of relatively few nuclear loci (25).

**Figure 1.**
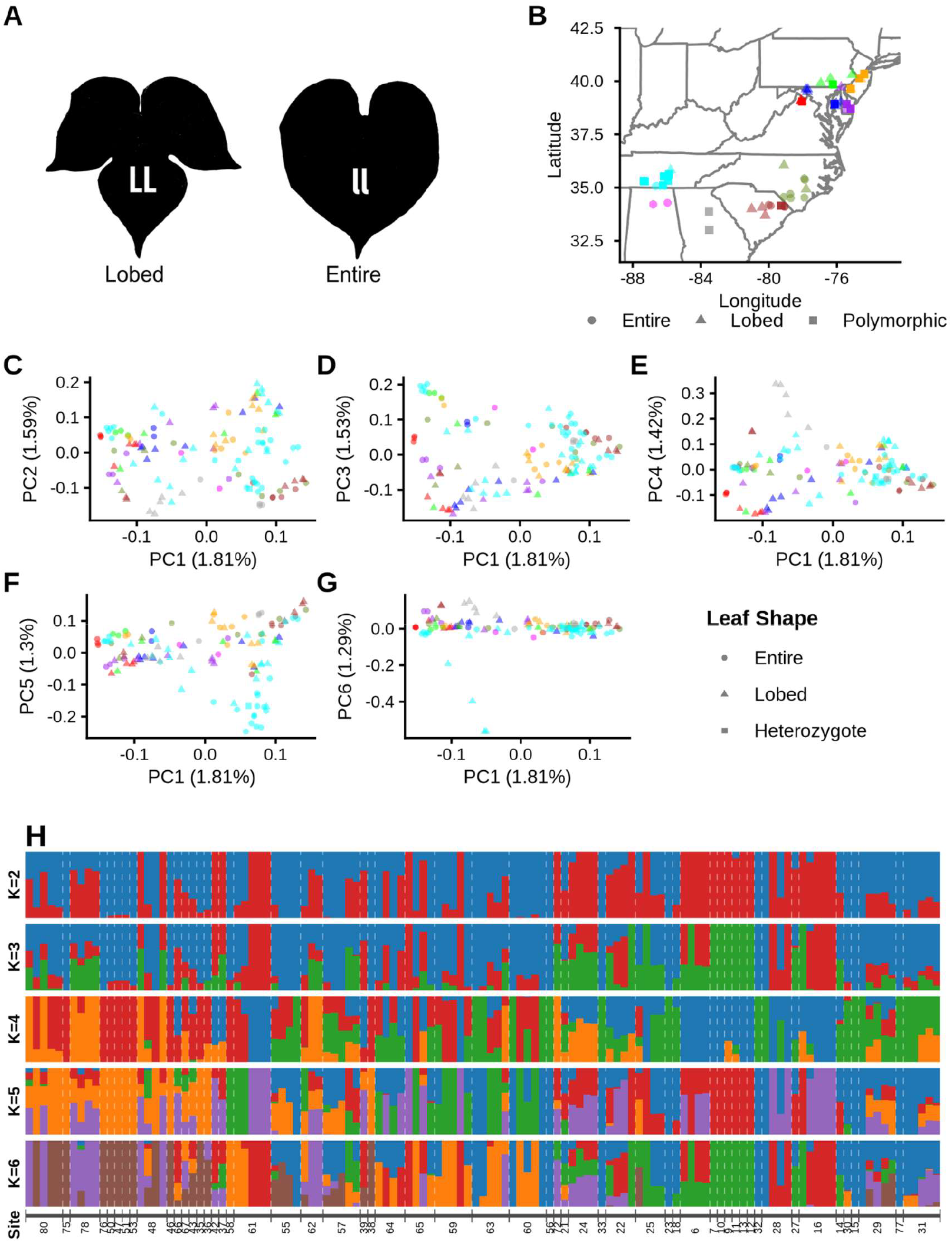
**A)** Illustrations of lobed (LL) and entire (ll) shaped leaves. **B)** Map of sequenced *I. hederacea* populations. Populations fixed for entire-shaped individuals are circles (N=1), populations fixed for lobed individuals are triangles (N=1), and polymorphic populations are squares (N=3 to 5). Colors represent sampling location, USA State. **C-G)** PC2-6 plotted against PC1 from PCA of neutral genetic diversity. Entire-shaped individuals are plotted as circles, lobed individuals are triangles, and the heterozygote individual is a square. Colors represent sampling location, USA State, as shown in A. **H)** Neutral genetic diversity ADMIXTURE plots of K2-6 with populations ordered by latitude. Individuals from different populations are separated by white dashed lines and the population sampling sites are denoted underneath the structure plots.

We used a combination of population genomics, genome-wide association studies (GWAS), and comparative genetics to characterise the genetic basis of leaf shape in *I. hederacea* and deduce the evolutionary history of the locus based on other *Ipomoea* species. To do so, we first assembled a chromosome-level *I. hederacea* reference genome and annotation. We then generated whole-genome sequencing for 123 individuals from 55 populations across the North American species range, finding weak signals of neutral population structure. We next conducted a GWAS and coverage association analysis that mapped the leaf shape locus to a 117 kb indel that perfectly co-segregates with the lobed phenotype. We then used a comparative genomic approach to show that the indel polymorphism is likely a newly derived deletion not present in five other closely related *Ipomoea* species.

## Results

### Chromosome-level reference genome assembly and annotation

We assembled a high quality chromosome-level haploid reference genome using PacBio long read sequencing from a highly homozygous lobed *I. hederacea* individual, VA12. The single-haplotype reference assembly size is 771.55 Mb, which is consistent with other closely related *Ipomoea* species; the reference assembly N50 is 42.53 Mb and L50 is 9, which compares favorably to other *Ipomoea* species. Contigs were anchored onto 15 chromosomes, 12 of which consisted of a single contig. The annotation of the repeat-masked reference assembly had a BUSCO completeness score of 99.6%, with 89.7% being single-copy genes (Table S1).

### Lack of neutral population structure

We first conducted a Principal Component Analysis (PCA) and ADMIXTURE v1.3.0 (30) for non-coding and synonymous single nucleotide polymorphisms (SNPs) to re-evaluate neutral population structure in *I. hederacea* using whole genome sequencing. Overall, we found no evidence of neutral population structure. We ran ADMIXTURE for K=1 to K=10 (Figure 1H; S2A-B) and K=1 is the best K-value with the lowest cross validation error (Figure S2C). For all K values, we found no evidence of clusters separated by either latitude or longitude (Figure 1H; S2A-B). The first 20 PCs explained a significant amount of neutral genetic variation according to a Tracy-Widom statistical test (Dataset S1). However, the first 20 PCs only explained between 0.94-1.81% each, and in total PC1-20 explained 23.27% of neutral genetic variation (Dataset S1). We observed very weak clustering when we plotted the PCs explaining the most amount of variation against each other and PCs did not clearly separate out populations sampled from the same region (Figure 1C-G). Although there was very weak clustering observed in PC space, we did find that PC2 and PC7 were significantly associated with latitude (Pearson correlation with Bonferroni correction: r_PC2_(121) = 0.56, *p*_*adj-PC2*_ < 0.001; r_PC7_(121) = −0.3, *p*_*adj-PC7*_ = 0.048; Figure S3). Similarly, PC5 was significantly associated with longitude (Pearson correlation with Bonferroni correction: r_PC5_(121) = 0.38, *p*_*adj-PC5*_ < 0.001; Figure S4). Given that leaf shape is geographically distributed, we also found that PC3 is significantly associated with leaf shape (Welch two sample t-test with Bonferroni correction: t_PC3_(119.58) = 9.89, *p*_*adj-PC3*_ < 0.001; Table S4). We provide further discussion of how the population genetic results support previous findings of an overall lack of neutral population genetic structure in *I. hederacea* in the supplementary material.

### A 117 kb indel perfectly co-segregates with leaf shape

To identify the leaf shape locus, we conducted a GWAS using GEMMA v0.98.5 (31). We quantified leaf shape for the GWAS by assigning entire-shaped seed families a numerical value of 0 and assigning lobed seed families a numerical value of 1. We found a 4.4 Mb region that was significantly associated with leaf shape on Chromosome 2 (Chr2:1164789-5609567). Considering that leaf lobing in *I. hederacea* is a Mendelian trait governed by a single locus, we expected an allele frequency difference between lobed and entire-shaped individuals to approach an absolute value of one. The two most significant SNPs in the leaf shape GWAS had an allele frequency difference of 0.98 (Chr2:3788962) and 0.93 (Chr2:3496407) between entire and lobed individuals (Figure 2C and Figure S5A). Both the GWAS and allele frequency differences suggest that the leaf shape locus is in close proximity to these two SNPs. However, we also observed low SNP density between the two leading GWAS SNPs, suggesting the presence of a structural variant (Figure S5C-D).

**Figure 2.**
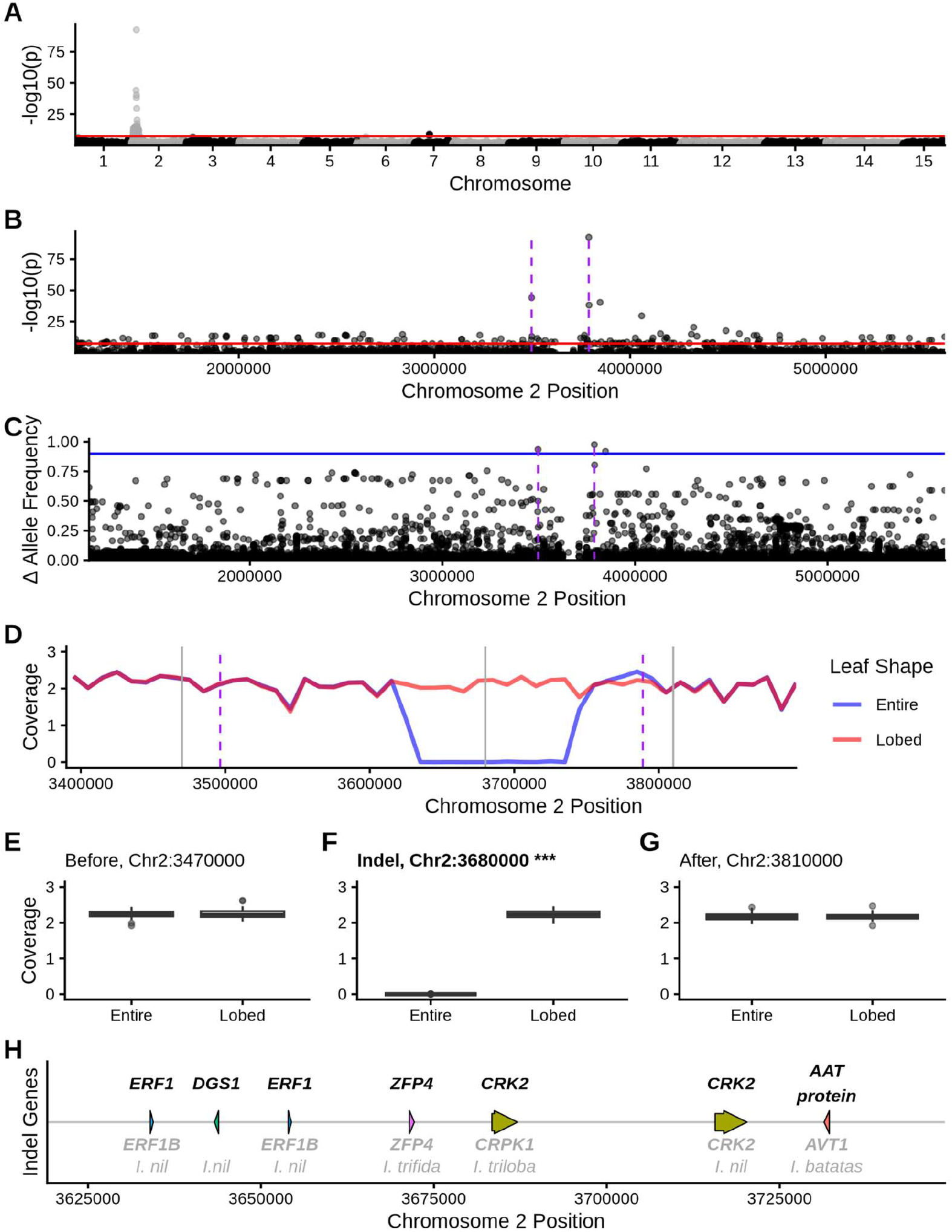
**A)** Leaf shape GWAS manhattan plot for the whole genome. **B)** Manhattan plot for peak on Chromosome 2. **A-B)** Red horizontal line is bonferroni corrected significance threshold. **C)** Allele frequency differences between individuals grouped by leaf shape within Chromosome 2 GWAS peak. The blue horizontal line shows the allele frequency difference of 0.9. **D)** Mean normalized coverage of entire (blue) and lobed (red) leaf shapes within the region surrounding leading GWAS SNPs. **B-D)** Purple dashed lines are the loci of the two SNPs with the highest GWAS significance and allele frequency difference. **E-G)** Boxplots comparing mean normalized coverage between leaf shape **E)** before the indel (Welch two sample t-test: t(118.36) = 0.28, *p* = 0.78), **F)** within the indel (Welch two sample t-test: t(60.033) = −173.8, *p* < 0.001), and **G)** after the indel (Welch two sample t-test: t(115.56) = −0.50, *p* = 0.62). **H)** Gene track with top BLAST hits when BLASTed against *A. thaliana* in black above each gene and top BLAST hit when BLASTed against the entire NCBI database in grey below each gene. Top BLAST hits that were not gene specific (e.g. gene family level or uncharacterized protein) are not included in the gene track without a label.

We next examined coverage differences between lobed and entire-shaped individuals to identify any structural variants that could be co-segregating with leaf lobing. We found a 117 kb region (Chr2:3625421-3742102) where entire-shaped individuals had a normalized mean coverage of ~0x but lobed individuals maintained a normalized coverage of ~2x which suggests the presence of an indel that perfectly co-segregates with leaf lobing (Figure 2D and S5B). Coverage significantly differed between entire and lobed individuals at a locus within the putative indel but not at two loci on either side of the indel (Welch two sample t-test: t_before_(118.36) = 0.28, *p*_*before*_ = 0.78; t_indel_(60.033) = −173.8, *p*_*indel*_ < 0.001; t_after_(115.56) = −0.50, *p*_*after*_ = 0.62; Figure 2E-G). These data suggest the large indel perfectly co-segregates with leaf shape in *I. hederacea*.

We next confirmed that the 117 kb region is actually a single deletion in the entire-shaped genotypes, rather than entire genotypes also possessing a structural variant missing from the lobed reference. To do so, we created a pseudo-reference genome of Chromosome 2 with the putative indel region removed, and realigned the short read sequencing to the pseudo-reference genome. For lobed genotypes that have genetic material present in the 117 kb region that is now absent in the pseudo-reference genome, we would expect short-read sequencing to no longer continuously align across the site where the indel was removed resulting in a gap in the alignment. If the entire-shaped genotypes have a single 117 kb deletion, we would expect reads to align continuously across the indel removal site in the pseudo-reference genome. As expected, lobed individuals had almost no reads aligning across the indel removal site with a mean of 0.11 reads per individual.

Entire-shaped individuals had a mean of 20.8 reads that continuously spanned the indel removal site, which is comparable to the number of reads before (mean = 21.61) and after (mean = 26.54) the indel region (Figure S10). These results indicate that the entire-shaped haplotype is likely a deletion rather than a more complex structural variant.

### Candidate genes within close proximity to putative indel

To identify candidate leaf shape genes on or near the indel, we BLASTed the 250 kb region on either side of the two most significant GWAS SNPs against the entire NCBI database as well as the *Arabidopsis thaliana* genome annotation (32). There are a total of 64 genes in the 793 kb region (Dataset S4-S5). There are seven genes in the putative leaf shape indel including two *A. thaliana* orthologs of *Ethylene Response Factor 1* (*ERF1*), one ortholog of *dgd1 suppressor 1* (*DGS1*), one ortholog of *zinc finger protein 4* (*ZFP4*), two orthologs of *cysteine-rich receptor-like protein kinase 2* (*CRK2*), and a transmembrane amino acid transporter family protein (Figure 2H, Table S2). We found similar results for orthologs in closely related *Ipomoea* species when we BLASTed the indel genes against the entire NCBI database (Figure 2H, Table S2). However, none of the genes on the indel are known to be directly involved in leaf shape or auxin biosynthesis. There are also two genes that are known to affect auxin distribution located within the 250 kb region of the leading GWAS SNPs: *Transmembrane Kinase 1* (*TMK1*) and *KANADI1* (*KAN1*; Table S3). The *TMK1* gene is 17 kb away from the second most significant SNP and 107 kb away from the indel and has been shown to be associated with leaf lobing in sweet potatoes (*I. batatas*) (17). *KAN1* is located 180 kb away from the indel and 133 kb from the most significant GWAS SNP and has been shown to affect leaf morphology in *I. nil* (33).

### Other leaf shape modifiers in indel region affect leaf shape in lobed genotypes

We next compared leaf shape genotypes for a variety of other traits that could be related to leaf shape, leaf morphology, size, and ecophysiology, to determine whether the indel polymorphism described above affected multiple leaf traits simultaneously, or mainly affected shape *per se*. We found that the two leaf shape genotypes did not differ in number of leaves, main vine length, stomatal conductance, leaf temperature, relative water content, trichome density, leaf area, leaf aspect ratio, and leaf roundness, as assessed in a greenhouse common garden experiment. However, we found a difference in leaf perimeter and leaf circularity, as expected, where lobed genotypes produce leaves with lower circularity and larger perimeters compared to entire genotypes (Welch two-sample t-test: t_perimeter_(121.88) = −8.84, *p*_*adj-perimeter*_ < 0.001; t_circularity_(81.91) = 48.67, *p*_*adj-circularity*_ < 0.001; Figure S6). These data suggest the indel affects shape *per se*, rather than other associated traits with secondary and more visible effects on shape.

A GWAS for leaf circularity with both lobed and entire-shaped individuals revealed a highly significant peak on Chromosome 2 in the same region as the leaf lobing GWAS, suggesting that loci affecting continuous indicators of shape are in the same region as the qualitative indicators we mapped above for lobing. We identified loci on Chromosomes 2, 6, and 7 that surpassed the Bonferroni corrected significance threshold for leaf circularity. Of the 127 SNPs significantly associated with leaf circularity, only one SNP is located on Chromosome 6, four SNPs are on Chromosome 7, and the remaining 122 SNPs are on Chromosome 2. Similar to the binary leaf shape GWAS above, the GWAS peak on Chromosome 2 was highly significant and spanned 4.44 Mb (Chr2:1164789 - 5609567; Figure S7A-B). The leaf circularity GWAS peak overlapped with the putative leaf shape indel and two most significant SNPs (Chr2:3788962 and Chr2:3496407) were the same SNPs in the lobing GWAS (Figure S7B).

We next hypothesized that any loci affecting leaf shape and development might have quantitative effects on shape *within* the lobed genotypes alone. A GWAS for leaf circularity using only lobed genotypes (N = 61) identified loci affecting subtle differences in shape variation. We detected a total of 25 loci on Chromosomes 2, 7, and 14 that surpassed the FDR 0.05 significance threshold but not a more stringent Bonferroni correction (Figure S7C). Similar to the leaf circularity GWAS for both lobed and entire individuals, the most significant peak was on Chromosome 2 which included 22 significant SNPs across 1.03 Mb (Chr2:3267701-4302532) that overlapped with the putative leaf shape indel (Figure S7D). However, the most significant SNP (Chr2: 3362671) was found 263 kb before the putative leaf shape indel (Figure S7D). We similarly performed a GWAS for only entire-shaped genotypes and found no SNPs that were significantly associated with leaf circularity (Figure S7E). Together, these results suggest that a locus in the region of Chromosome 2 affects leaf shape both within and between alternative leaf shape genotypes.

### Indel is a newly derived deletion in *I*. *hederacea*

We next tested for the presence of the indel in other closely related *Ipomoea* species with publicly available genomic data. We used five *Ipomoea* species that have reference genomes and also happen to have different leaf shapes. To eliminate potential differences from genome annotation pipelines, we re-annotated genomes for *Ipomoea* species: *I. aquatica* (34); *I. cairica* (35); *I. nil* (version Asagao1.2 downloaded from http://viewer.shigen.info/asagao/download.php) (36); *I. purpurea* (37); *I. triloba* (38). We then BLASTed the *I. hederacea* indel genes against the other *Ipomoea* species and used GENESPACE to identify syntenic orthologs. We detected an extensive amount of genome-wide synteny between the six *Ipomoea* species (Figure 3A; S8). More importantly, we found syntenic orthologs for multiple of the genes on the leaf shape indel across the five other *Ipomoea* species regardless of leaf shape (Figure 3B, Table S5), suggesting that the indel is a newly arisen deletion in *I. hederacea*.

**Figure 3.**
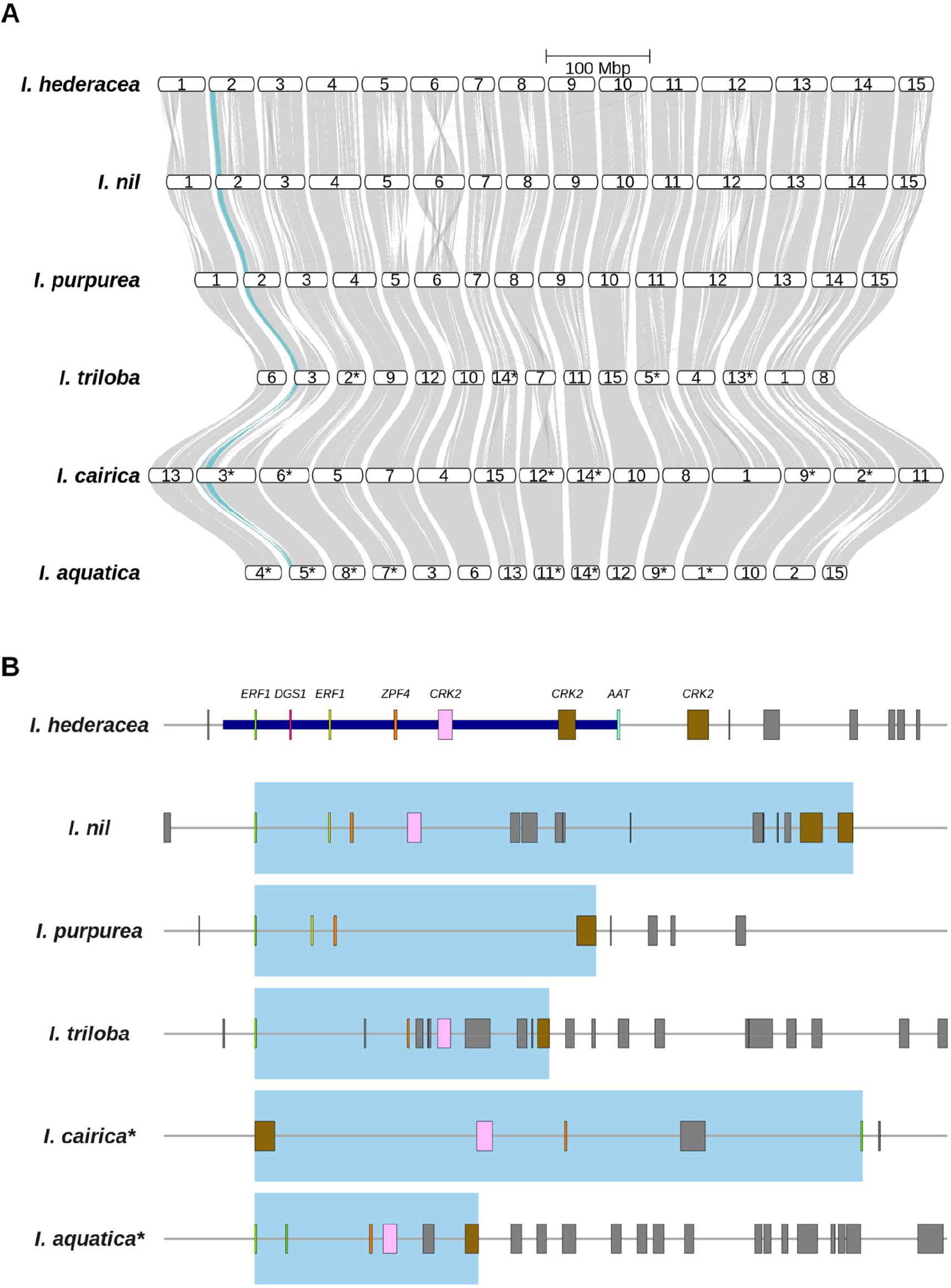
**A)** Genome-wide riparian plot across six *Ipomoea* species. The syntenic region for the leaf shape GWAS peak on Chromosome 2 in *I. hederacea* is highlighted in blue. **B)** Syntenic orthologs of indel genes in other *Ipomoea* species. The thick dark blue line indicates the indel region in *I. hederacea*. The blue shaded boxes in other *Ipomoea* species show the regions containing syntenic orthologs the indel gene. **A-B)** Chromosomes and regions denoted with an asterisk next to the species name have been inverted to match the orientation of the *I. hederacea* reference genome assembly (Figure S10).

Most species, except for *I. triloba*, have two ERF genes (g17358 and g17360) in the syntenic region, although not all of the ERF genes belong to the same orthogroup. The first *I. hederacea* ERF gene on the indel has a syntenic ortholog for all *Ipomoea species* and *I. aquatica* has a tandem duplication for this gene. The other ERF gene found on the *I. hederacea* leaf-shape indel only has a syntenic ortholog in *I. nil* and *I. purpurea*. However, *I. cairica* has another ERF gene located in the same syntenic block that does not belong to the same orthogroup as the other ERF genes found in the other species. Similarly, from BLASTing the *I. hederacea* indel genes against the other species, we found evidence that the one ERF gene in the indel is found across all species and the second *I. hederacea* ERF gene only has unique BLAST hits in *I. nil* and *I. purpurea* (Table S5).

There are varying numbers of CRK genes found in the leaf-shape indel across the different *Ipomoea* species and not all the CRK genes are from the same orthogroup. In *I. hederacea*, there are two CRK genes on the indel (Figure 2H, Table S2). According to both GENESPACE and BLAST, the first CRK *I. hederacea* gene (g17362) has a single syntenic hit in all species except for *I. purpurea* (Table S5; Dataset S2). However, all species have at least one syntenic ortholog for the second CRK gene found on the indel (g17363; Figure 3B; Table S5). Both *I. hederacea* and *I. nil* have two copies of the second CRK, but only one of them is actually on the indel in *I. hederacea* whereas the other copy is located next to the indel (g17365; Figure 3B). Additionally, there is another CRK orthogroup present in the same syntenic region in other *Ipomoea* species that is missing in *I. hederacea* where *I. aquatica* has a single copy but *I. triloba* and *I. nil* have two copies (Dataset S2). Overall, we observe variation in the presence and number of CRK orthologs between *Ipomoea* species, but we do not find any evidence that these differences correspond to leaf shape differences between the species.

Both the GENESPACE and BLAST results suggest that there were no syntenic orthologs for the *I. hederacea* dgd1 suppressor gene (g17359) or the transmembrane AA transporter gene (g17364) in the other *Ipomoea* species (Table S5; Dataset S2).

Despite *I. purpurea* being the most closely related species with predominantly entire-shaped leaves, we found no evidence of the deletion segregating in 78 *I. purpurea* individuals that were previously sequenced by Gupta et al. (37). We aligned whole genome sequencing from 78 *I. purpurea* individuals to the *I. hederacea* reference genome to determine if there were reads aligning to the putative leaf shape indel region. If the indel was segregating in the *I. purpurea* samples, we would expect to see some individuals with consistently ~0x coverage across the indel region similar to entire-shaped *I. hederacea* individuals. All *I. purpurea* individuals had reads aligning within the indel region suggesting that the 117 kb deletion is not a genetic variant present in any of the sequenced *I. purpurea* individuals (Figure S9). However, the coverage across the indel region is not consistently at ~2x normalized coverage as seen in the *I. hederacea* lobed individuals, which is likely as a result of aligning the *I. purpurea* reads to the *I. hederacea* reference genome. From visually inspecting a small subset of bam files in IGV v2.18.4 (39, 40), the drops in coverage appear in close proximity with genes that are missing syntenic orthologs in *I. purpurea* (Figure S9, Table S5, Dataset S2).

These results suggest that the 117 kb deletion associated with the entire shape phenotype in *I. hederacea* is not present in *I. purpurea* despite the similarity in leaf shape phenotype.

## Discussion

We identified a 117 kb indel in *I. hederacea* that perfectly co-segregates with the lobed leaf shape phenotype. The presence of syntenic orthologs in closely related *Ipomoea* species for the genes within the putative indel indicates that it is a newly arisen polymorphism in *I. hederacea*. The mapping and comparative genomics data indicate that the repeated evolution of similar leaf shape phenotypes in congeneric *Ipomoea* species is likely governed by different genetic mechanisms.

Below, we discuss potential mechanisms for how the indel could be affecting leaf shape, pleiotropic effects of the indel, leaf shape in the *Ipomoea* genus, and how the discovery of the indel more broadly informs our understanding about the ecology and evolution of leaf shape variation.

### Indel regulating gene expression of candidate genes

Structural variants, such as large insertions-deletions (indels), have proven to be an important component of genetic variation and have been linked to multiple phenotypic traits including leaf shape variation in other species (41). Indels can have widespread phenotypic effects as a consequence of one haplotype missing important gene content that can range from portions of a single gene to missing entire genes (42–44). The lack of recombination between alternative indel haplotypes can cause adjacent genes to be inherited together, sometimes referred to as a supergene, which can impact multiple traits simultaneously (45). The phenotypic effects of indels are not only as a result of the absence or presence of genes in the indel, but structural variants can also affect the gene expression of other genes by introducing new regulatory elements (46), changing distances between regulatory elements (47, 48), and affecting 3D configurations of DNA (48).

Although indels have been shown to be mostly under purifying selection some indels are still maintained in natural populations (43, 44, 46, 49), can facilitate local adaptation (50), are correlated with environmental factors (42, 49), and can directly impact fitness (42, 50).

Although the indel itself does not contain known candidate genes with annotations implicating leaf shape, three pieces of evidence suggests it affects the expression of other genes involved in leaf shape and development. First, the GWAS on leaf circularity using only lobed genotypes detected significant SNPs affecting continuous shape variation, in the same region as the indel (Figure S7C-D). Second, a bulked segregant analysis examining leaf shape variation within the *I. hederacea* lobed genetic background similarly finds a significant QTL region that overlaps with the indel (9).

Third, linkage mapping in *I*.*nil* mapped multiple leaf shape genes to the tip of Chromosome 2 (19–22, 36). In *I. hederacea*, there are possible candidate genes located within 250 kb of the two most significant GWAS SNPs including orthologs for *TMK1*, and *KAN1. TMK1* is involved in the auxin synthesis pathway and has been shown to bind and phosphorylate PIN proteins that affect auxin concentration in the cell (51). Auxin is a plant hormone that is known to affect plant growth, and more specifically has been directly linked to leaf shape variation (52, 53). The *TMK1* gene is located between two leading GWAS SNPs 17 kb away from the second most significant GWAS SNP and 107 kb away from the indel. The *TMK1* ortholog is the most promising candidate gene given its close proximity to the most significant genetic variants and that it has also been identified as a leaf shape candidate gene in *I. batatas* (17). *KAN1* is also involved in the auxin synthesis pathway and affects the development of the lower leaf surface (54, 55). An ortholog of *KAN1*, known as the *FEATHERED* gene, affects leaf morphology in *I. nil* (33). To rigorously test if the indel is affecting leaf shape by regulating the expression of other genes would require developing near-isogenic lines differing only at the leaf shape locus to test for differences in expression in the same genetic background across a series of developmental stages. Note that comparing expression among naturally occurring inbred lines would likely identify expression differences unrelated to leaf shape as flowering time, growth rate, and flower morphology also show latitudinal clines across the same range (23, 24). Alternatively, since the genetic basis of leaf shape is known to differ between plant taxa (1), it is also plausible that one of the indel genes could be directly impacting leaf shape *I. hederacea* but orthologs in other species that have been functionally validated do not have the same effect on leaf shape. Although the proximate mechanisms by which this region affects leaf shape require further investigation (and ideally genetic manipulation), the results indicate that the indel is the leaf shape locus, as it is the only genetic variant to perfectly co-segregate with the leaf shape phenotype.

### Possible pleiotropic effects of the indel on multiple traits

Differences in coverage between lobed and entire-shaped individuals suggest that the indel perfectly co-segregates with the leaf shape phenotypes where all entire-shaped individuals are missing the 117 kb region. There are seven genes in the indel including *A. thaliana* orthologs for *ERF1, DGS1, ZFP4, CRK2*, and an amino acid transporter family protein. While none of these genes are particularly strong candidates for affecting leaf shape, their annotations and functions suggest that their deletion could affect other ecologically important traits. For example, *ERF1* is involved in the ethylene signaling pathway, jasmonic acid pathway, and has been linked to drought and stress tolerance as a consequence of differences in stomatal aperture (56–58). *DGS1* is involved in the galactolipid biosynthesis pathway and hydrogen peroxide biosynthesis pathway and is associated with drought response (59, 60). *CRK2* has been shown to be associated with defense against pathogens (61). Collectively, the deletion of these genes could explain some of the other phenotypic traits that are associated with leaf shape in *I. hederacea* such as herbivory preference (28), pathogen resistance (3), microbial community (29), and stomatal conductance in the field (27). The differences in plant defense and physiology that have been observed between the leaf shape genotypes (3, 27–29) thus might not be due to differences in leaf shape itself but rather the pleiotropic effects of multiple genes present on the indel. Therefore, it is possible that the latitudinal leaf shape cline might be driven by the combined effect of multiple beneficial traits associated with the indel (see supplementary material).

### Indel is a newly arisen deletion in *I*. *hederacea*

Despite the prevalence of similar leaf shape phenotypes across the *Ipomoea* genus, we show that the entire-shape haplotype is a new deletion in *I. hederacea*. Firstly, we determined that the entire-shape haplotype is likely a deletion, as opposed to a more complex structural variant, given that sequencing reads for the entire-shape genotype span across both sides of the indel after removing the indel from the reference genome (Figure S10). Secondly, we found syntenic orthologs for genes on the putative leaf shape indel in the five other *Ipomoea* species (Figure 3B). Although we observed variation in the presence and number of orthologs for some indel genes between species, we did not find any evidence that this variation was associated with leaf shape differences between the species. Even *I. purpurea*, a close relative with nearly identical entire-shaped leaves to *I. hederacea*, has the indel region present in 78 genetic lines previously sequenced by Gupta et al. (2023, Figure 3B, Figure S9). Given these data suggesting that the indel polymorphism is specific to *I. hederacea*, there must be other genetic mechanisms giving rise to similar leaf shape phenotypes across the *Ipomoea* genus. Early linkage mapping studies of ornamental *I. nil* cultivars indicated that there are indeed multiple genes in several different linkage groups affecting leaf shape (19–22). Collectively, we find evidence that the leaf shape genetic mechanism we characterized is a newly arisen deletion in *I. hederacea* that has resulted in the repeated evolution of similar leaf shape phenotypes in the *Ipomoea* genus via a unique genetic mechanism.

### Future directions

Our work shows that a simple Mendelian polymorphism that exhibits a clinal distribution, likely due to selection, is determined by a 117 kb indel polymorphism that is newly arisen in *I. hederacea*. Further, the indel polymorphism for leaf shape is not present in related species with similar leaf shapes, indicating that similar leaf morphology can have independent genetic mechanisms even in closely related species. The observation that the indel includes multiple genes involved in plant defense, ecophysiology, and plant-pathogen interactions suggests that the ecological mechanisms underlying clinal variation in *I. hederacea* could be a consequence of the correlated effects of the indel on multiple genes and traits that are inherited together, rather than any single ecological mechanism acting in isolation. These data, and developing genomic resources for *Ipomoea* (14, 16, 17, 34–38, 62), set the stage for detailed investigations of the genetic, developmental, and ecological mechanisms governing leaf shape variation within the genus.

## Materials and Methods

### Sampling design and phenotyping

We used a subset of seed families collected by Campitelli and Stinchcombe (2013) (4). We randomly selected one seed family from each population that was fixed for either entire or lobed leaf shape. For polymorphic populations, where possible we selected up to five seed families (two entire, two lobed, and one heterozygote). If there were not enough seed families of a given leaf shape in a polymorphic population, we then selected another seed family from one of the remaining leaf shapes. Our final sample included 123 seed families (61 lobed, 61 entire, and one heterozygote seed family) from 55 populations (22 lobed, 13 entire, and 20 polymorphic populations; Figure 1B). For each seed family, we collected phenotypic data in a greenhouse common garden experiment on leaf shape (entire vs lobed, area, circularity, roundness, perimeter, aspect ratio), leaf counts, leading vine length, trichome density, relative water content, leaf surface temperature, and stomatal conductance (supplementary methods).

### Genome assembly, resequencing, and variant calling

We chose a lobed seed family, VA12, from a polymorphic population in the middle of the range for the reference genome assembly. A chromosome-scale reference genome was generated using PacBio HiFi long-reads and Dovetail Omni-C for scaffolding (supplementary methods). Contigs were manually ordered and re-oriented based on the contact map, and chromosomes were renamed against the *Ipomoea nil* genome in juicer_tools v1.22.01 (63) and juicebox v2.20.00 (64). We annotated the reference genome using homology based evidence from OrthoDB v. 11 Viridiplatae proteins downloaded from https://bioinf.uni-greifswald.de and RNA short-read sequencing of different tissue types (supplementary methods).

We also generated whole-genome short-read resequencing for the 123 seed families, using Illumina TruSeq PCR-free libraries sequenced on an Illumina NovaSeq X with PE150 reads (supplementary methods). We aligned the short-read sequencing for all 123 seed families to the reference genome to create a VCF file filtered for quality, missingness, and depth (supplementary methods). The VCF used in all downstream analyses had a total of 2,990,321 biallelic SNPs after filtering.

### Population structure

We used 4-fold degenerate protein-coding sites and non-coding sites to evaluate neutral genetic diversity. We determined degeneracy of protein coding sites using the codingSiteTypes.py script from https://github.com/simonhmartin/genomics_general. We then removed all protein coding sites that were not 4-fold degenerate sites from the biallelic SNP VCF. Given the differences observed in lower minor allele frequencies between the northern and southern parts of the range, we decided to only remove singletons and homozygous doubletons because lower minor allele frequencies could be informative for *I. hederacea* population structure (65). We used vcftools v0.1.16 (66) to identify and remove singletons and homozygous doubletons from the biallelic SNPs VCF. To identify an appropriate linkage disequilibrium (LD) pruning threshold, we estimated average linkage disequilibrium decay (Figure S1, supplementary methods).

To examine neutral population structure, we conducted a Principal Component Analysis (PCA) and used ADMIXTURE. We generated 122 Principal Components (PCs) in PLINK v1.9b_6.21-x86_64 (67). For all follow up analyses we used the first 20 Principal Components (PCs) that explained a significant amount of variation according to a Tracy-Widom test from R package AssocTests v 1.0-1 (68). We calculated the Pearson correlation between PCs and latitude or longitude to determine if PCs were associated with geography. We also used Welch’s two sample t-tests to determine if any PCs are significantly associated with leaf shape. We used a Bonferroni correction to adjust p-values (*p*_*adj*_ = *p*_*raw*_*60) to account for multiple comparison testing of PCs against latitude, longitude, and leaf shape. We ran ADMIXTURE v1.3.0 (30) with K values 1-10 to determine if there was any neutral population structure. We used the lowest cross validation error to determine the best K-value.

Structure plots were visualized using R package pophelper v2.3.1 (69).

### Identifying the lobed locus

We conducted a GWAS and a coverage association analysis to identify genetic variants that are associated with leaf shape. For the GWAS, we used GEMMA v0.98.5 to identify SNPs associated with leaf shape (31). We used bcftools view v1.19 (70) to exclude the heterozygote seed family from the biallelic SNP VCF file and to sort samples in the same order as the phenotype input file. We used beagle v5.4-240301 to impute missing genotypes (71). Bcftools annotate was used to create unique SNP IDs for all variants. GEMMA genotype, annotation, and phenotype input files were formatted using a custom bash script. For the genotype input file, homozygotes for the reference allele were assigned a value of 0, heterozygotes were assigned a value of 1, and homozygotes for the alternative allele were assigned a value of 2. We used the GEMMA univariate linear mixed model algorithm with leaf shape as a quantitative trait by assigning a numerical value of 0 to entire-shaped seed families and 1 to lobed seed families. We used GEMMA to estimate a centered relatedness matrix used to correct for population structure in the GWAS. The GWAS included 1,293,141 SNPs after removing SNPs with a minor allele frequency < 5% or SNPs that had a correlation with covariates > 0.9999.

We would expect that alternative alleles will be fixed between lobed and entire-shaped individuals because leaf shape is a Mendelian trait. Therefore, we calculated the allele frequency difference between lobed and entire-shaped individuals with the expectation that the leaf shape locus should have an allele frequency difference approaching one. We used bcftools view to generate two separate VCF files for the different leaf shapes that only included biallelic SNPs from Chromosome 2. We then used the bcftools plugin (INFO/AF) to calculate the allele frequency at each locus and bcftools query to create tab-separated text files that had the allele frequency for each locus. We calculated and visualised allele frequency differences between leaf shapes in R v4.4.0 (72).

For the coverage association analysis, we calculated the mean coverage in 10 kb windows across Chromosome 2 from aligned bam files using mosdepth v0.3.10 (73). By default, mosdepth excludes all reads previously flagged by samtools as being unmapped, not a primary alignment, failing quality checks, or a read duplicate. We additionally filtered out reads that had a mapping quality score less than 30 that is inline with the mapping quality score cutoff we used for calling SNPs. To compare coverage between leaf shapes, we calculated the mean normalized coverage for both leaf shapes. To calculate the normalized mean coverage for each seed family, we divided the coverage of each 10 kb window by the mean coverage of Chromosome 2 for a given seed family and then multiplied by two because *I. hederacea* is a diploid. We conducted a t-test comparing normalized coverage between leaf shapes at a locus ~25 kb on either side of the two most significant GEMMA SNPs (Chr2:3470000 and Chr2:3810000) as well as a locus (Chr2:3680000) within a potential indel located in a region of low SNP density between the two most significant GWAS SNPs (Figure S5C-D).

We BLASTed genes to infer gene function and identify candidate genes. Given that the Chromosome 2 region that surpasses the Bonferroni threshold is ~4.44 Mb, we chose to prioritize a narrower region within the peak that is most significant when looking for candidate genes. We used gffread v0.12.3 (74) and SeqKit v2.5.1 (75) to generate a list of genes found in a 793 kb region (Chr2:3246407-4038962) that spans 250 kb on either side of the two most significant GWAS SNPs (Chr2:3496407 and Chr2:3788962). We used BLASTp v2.14.1 (76) to BLAST the 793 kb region against the entire NCBI database and an *A. thaliana* protein database that we made from the *A. thaliana* TAIR10 annotation (32) downloaded from NCBI.

### Leaf circularity GWAS

For leaf circularity, we conducted three follow up GWAS analyses. The first GWAS included all lobed and entire-shaped seed families (n = 122). We then conducted a GWAS with only lobed seed families (n = 61) and a separate GWAS with only entire-shaped seed families (n = 61). The GWAS that included both lobed and entire-shaped seed families was using the same VCF and GEMMA parameters as described in the “Identifying the lobed locus” section. We regenerated VCFs for the GWAS with only lobed or only entire-shaped seed families because variants that are specific to a leaf shape phenotype (e.g. SNPs present on the indel region) might be filtered out of the VCF file with all individuals. To do so, we filtered a VCF file with all 123 seed families that included both variant and invariant sites. We used bcftools to keep only lobed or only entire-shaped seed families and bi-allielic SNPs as described in the “Aligning to reference genome and calling SNPs” section.

For all three univariate GWAS and generating all input files was done as described in the “Identifying the lobed locus” section. The GWAS with 122 seed families had 1,293,141 SNPs, the GWAS with only lobed individuals had 948,935 SNPs, and the GWAS with only entire-shaped individuals had 1,459,872 SNPs after removing SNPs with a minor allele frequency <5% or SNPs that had a correlation with covariates >0.9999.

### Comparative genomics

To determine if the leaf shape indel is conserved across *Ipomoea* species, we used previously published reference genomes of other *Ipomoea* species: *Ipomoea aquatica* (34), *Ipomoea cairica* (35), *Ipomoea nil* (version Asagao1.2 downloaded from http://viewer.shigen.info/asagao/download.php) (36), *Ipomoea purpurea* (37), *Ipomoea triloba* (38). To ensure that differences observed among species were not because of differences in annotation methods, we reannotated all genomes used in the comparative genomic analyses. The *I. purpurea* genome was repeat-masked with RepeatModeler v2.0.3 (77) and RepeatMasker v4.1.4 (78). Other genomes were already masked previously. The full length transcripts of *I. aquatica* and *I. cairica*, and the *I. purpurea* RNA-seq short reads were deduplicated with bbmap-dedupe (79). Other RNA-seq short reads were cleaned similar to *I. hederacea* as described in the ‘Genome assembly and annotation’ section. The unique RNA-seq reads or assembled contigs and the OrthoDB v.11 Viridiplantae proteins downloaded from https://bioinf.uni-greifswald.de were used as evidences in the genome annotation with BRAKER genome annotator (80). The *I. cairica* genome was annotated with the OrthoDB Viridiplantae proteins alone due to a low number of transcripts retained after running dedupe with qtrim was turned on.

To compare syntenic regions between genomes, we used GENESPACE v1.3.1 (81) with v2.1.8 (82), orthofinder v.2.5.4 (83, 84), and MCScan v1.0.0 (85). Input fasta and gff files were formatted using the genespace parse annotations function and then we further filtered for the longest isoform using the R package Biostrings v2.74.1 (86). GENESPACE was run using default parameters.

Orthologs within the leaf shape indel were identified by querying the GENESPACE pan-genomes annotation for Chr2:3625000-3755963. As a complementary approach, we created BLAST databases of the other *Ipomoea* species and BLASTed the indel region in *I. hederacea* against the other *Ipomoea* species as described in “Identifying the leaf shape locus” section. We used GENESPACE and ggplot2 v4.0.0 (87) to generate a genome-wide riparian plot to compare synteny across all six *Ipomoea* species. We used the synthetic hits output from GENESPACE to generate dot plots using ggplot2.

Given that the reference genome for each species only represents a single individual, we checked if there were reads mapping to the indel region in the closest related entire-shaped species, *I. purpurea*. We downloaded whole genome sequencing of 79 *I. purpurea* individuals that was previously sequenced by Gupta et al. (2023) (37) from the NCBI short-read archive using sra-toolkit v3.0.9 (88). We ran fastqc v.0.12.1 (89) and multiqc v.1.19 (90) to assess quality. We trimmed adapter sequences with a 3’ 5 bp clip using TrimGalore v0.6.10 (91) and excluded one individual with skewed GC content. We then aligned the 78 *I. purpurea* individuals to the *I. hederacea* reference genome and generated bam files as described in “Aligning to reference genome and calling SNPs” section. To check if reads were aligning to the indel region, we calculated the normalized mean coverage in 10 kb windows as described for the coverage association analysis in the “Identifying the lobed locus” section.

## Supporting information

Supplement including: supporting methods, supporting results, Figures S1-S11, Tables S1-S5, Legends for Datasets S1-S5

Dataset S1. Tracy-widom statistic for population genetics PCA

Dataset S2. Queried GENESPACE pangenomes file for indel region

Dataset S3. Phenotype data for homozygote seed families (n=122) measured in common garden experiment.

Dataset S4. Top BLAST hits for I. hederacea Chromosome 2 GWAS peak region BLASTed against A. thaliana

Dataset S5. Top BLAST hits for I. hederacea Chromosome 2 GWAS peak region BLASTed against NCBI nr database

## Acknowledgements

Joanna Rifkin, Stephen Wright, Aneil Agrawal, and members of the Stinchcombe provided valuable feedback on the project. We would like to thank Savannah Lollo, Ishani Sharma, Grace Walker Mitchell, Aidan Godin and Jessica Underwood for helping with the common garden experiments.

Katie Monat also helped with data entry and ImageJ processing. We are extremely grateful for all the help from William Cole, Thomas Gludovacz, and Alice DesRoches whilst using the UofT growth facilities. Our research was enabled by the computational resources and support provided from Compute Ontario and the Digital Research Alliance of Canada.

## Funding sources

NSF IOS-PGRP CAREER Award 2239530 to AH.

NSERC CGSM + PGSD to ALP.

NSERC discovery to JRS.

## References

1. A. B. Nicotra, et al., The evolution and functional significance of leaf shape in the angiosperms. Funct. Plant Biol. 38, 535–552 (2011).

2. R. Wyatt, J. Antonovics, Butterflyweed re-revisited: Spatial and temporal patterns of leaf shape variation in Asclepias Tuberosa. Evolution 35, 529–542 (1981).

3. K. L. Bright-Emlen, “Geographic variation and natural selection on a leaf shape polymorphism in the ivyleaf morning glory (Ipomoea hederacea),” Duke University, United States --North Carolina. (1998).

4. B. E. Campitelli, J. R. Stinchcombe, Natural selection maintains a single-locus leaf shape cline in Ivyleaf morning glory, Ipomoea hederacea. Mol. Ecol. 22, 552–564 (2013).

5. K. G. Ferris, et al., Leaf shape evolution has a similar genetic architecture in three edaphic specialists within the Mimulus guttatus species complex. Ann. Bot. 116, 213–223 (2015).

6. Z. Migicovsky, M. Li, D. H. Chitwood, S. Myles, Morphometrics reveals complex and heritable apple leaf shapes. Front. Plant Sci. 8, 2185 (2018).

7. P. Liu, et al., Enhanced genome-wide association reveals the role of YABBY11-NGATHA-LIKE1 in leaf serration development of Populus. Plant Physiol. 191, 1702–1718 (2023).

8. K. L. Bright, M. D. Rausher, Natural selection on a leaf-shape polymorphism in the Ivyleaf Morning Glory (Ipomoea hederacea). Evolution 62, 1978–1990 (2008).

9. M. Henry, “The genetic basis of the 5-lobed leaf shape in Ivyleaf Morning Glory (Ipomoea hederacea),” University of Toronto. (2025).

10. S. Andersson, Geographical variation and genetic analysis of leaf shape in Crepis tectorum (Asteraceae). Plant Syst. Evol. 178, 247–258 (1991).

11. B. A. Gould, J. R. Stinchcombe, Population genomic scans suggest novel genes underlie convergent flowering time evolution in the introduced range of Arabidopsis thaliana. Mol. Ecol. 26, 92–106 (2017).

12. C. A. Wessinger, L. C. Hileman, Parallelism in flower evolution and development. Annu. Rev. Ecol. Evol. Syst. 51, 387–408 (2020).

13. H. A. Orr, The probability of parallel evolution. Evolution 59, 216–220 (2005).

14. R. S. Baucom, S.-M. Chang, J. M. Kniskern, M. D. Rausher, J. R. Stinchcombe, Morning glory as a powerful model in ecological genomics: tracing adaptation through both natural and artificial selection. Heredity 107, 377–385 (2011).

15. A. Jones, J. Kang, Development of leaf lobing and vein pattern architecture in the genus Ipomoea (Morning Glory). Int. J. Plant Sci. 176, 820–831 (2015).

16. S. Gupta, D. M. Rosenthal, J. R. Stinchcombe, R. S. Baucom, The remarkable morphological diversity of leaf shape in sweet potato (Ipomoea batatas): The influence of genetics, environment, and G×E. New Phytol. 225, 2183–2195 (2020).

17. D. An, et al., Sweet potato gene clusters control anthocyanin biosynthesis and leaf morphology. Plant Biotechnol. J. 1–28 (2026). 10.1111/pbi.70636.

18. C. D. Elmore, Mode of reproduction and inheritance of leaf shape in Ipomoea hederacea. Weed Sci. 34, 391–395 (1986).

19. Y. Imai, A genetic monograph on the leaf form of Pharbitis Nil. Z. Für Indukt. Abstamm.- Vererbungslehre 55, 1–107 (1930).

20. Y. Imai, Description of the genes found in Pharbitis Nil. Genetica 12, 297–318 (1930).

21. Y. Imai, Linkage studies in Pharbitis Nil. III. Z. Für Indukt. Abstamm.- Vererbungslehre 66, 219–235 (1934).

22. Y. Imai, Linkage groups of the Japanese Morning Glory. Genetics 14, 223–255 (1929).

23. A. J. Stock, B. E. Campitelli, J. R. Stinchcombe, Quantitative genetic variance and multivariate clines in the Ivyleaf morning glory, Ipomoea hederacea. Philos. Trans. R. Soc. B Biol. Sci. 369, 20130259 (2014).

24. G. A. Henry, J. R. Stinchcombe, G-matrix stability in clinally diverging populations of an annual weed. Evolution 77, 49–62 (2023).

25. B. E. Campitelli, John. R. Stinchcombe, Population dynamics and evolutionary history of the weedy vine Ipomoea hederacea in North America. G3amp58 GenesGenomesGenetics 4, 1407–1416 (2014).

26. B. E. Campitelli, A. J. Gorton, K. L. Ostevik, J. R. Stinchcombe, The effect of leaf shape on the thermoregulation and frost tolerance of an annual vine, Ipomoea hederacea (Convolvulaceae). Am. J. Bot. 100, 2175–2182 (2013).

27. Y. K. Singhal, J. A. Boyle, J. R. Stinchcombe, Differences in stomatal conductance between leaf shape genotypes of Ipomoea hederacea suggest divergent ecophysiological strategies. MicroPublication Biol. (2025). 10.17912/micropub.biology.001528.

28. B. E. Campitelli, A. K. Simonsen, A. Rico Wolf, J. S. Manson, J. R. Stinchcombe, Leaf shape variation and herbivore consumption and performance: a case study with Ipomoea hederacea and three generalists. Arthropod-Plant Interact. 2, 9–19 (2008).

29. J. A. Boyle, M. E. Frederickson, J. R. Stinchcombe, Genetic architecture of heritable leaf microbes. Microbiol. Spectr. 12, e0061024 (2024).

30. D. H. Alexander, J. Novembre, K. Lange, Fast model-based estimation of ancestry in unrelated individuals. Genome Res. 19, 1655–1664 (2009).

31. X. Zhou, M. Stephens, Efficient multivariate linear mixed model algorithms for genome-wide association studies. Nat. Methods 11, 407–409 (2014).

32. D. Swarbreck, et al., The Arabidopsis Information Resource (TAIR): gene structure and function annotation. Nucleic Acids Res. 36, D1009–1014 (2008).

33. M. Iwasaki, E. Nitasaka, The FEATHERED gene is required for polarity establishment in lateral organs especially flowers of the Japanese morning glory (Ipomoea nil). Plant Mol. Biol. 62, 913–925 (2006).

34. F. Jiang, et al., Improved chromosome-level genome and annotation data for a leafy vegetable water spinach (Ipomoea aquatica). Sci. Hortic. 320, 112193 (2023).

35. F. Jiang, et al., A chromosome-level reference genome of a Convolvulaceae species Ipomoea cairica. G3 GenesGenomesGenetics 12, jkac187 (2022).

36. A. Hoshino, et al., Genome sequence and analysis of the Japanese morning glory Ipomoea nil. Nat. Commun. 7, 13295 (2016).

37. S. Gupta, et al., Interchromosomal linkage disequilibrium and linked fitness cost loci associated with selection for herbicide resistance. New Phytol. 238, 1263–1277 (2023).

38. S. Wu, et al., Genome sequences of two diploid wild relatives of cultivated sweetpotato reveal targets for genetic improvement. Nat. Commun. 9, 4580 (2018).

39. J. T. Robinson, et al., Integrative Genomics Viewer. Nat. Biotechnol. 29, 24–26 (2011).

40. H. Thorvaldsdóttir, J. T. Robinson, J. P. Mesirov, Integrative Genomics Viewer (IGV): high-performance genomics data visualization and exploration. Brief. Bioinform. 14, 178–192 (2013).

41. H. Bastiaanse, et al., A systems genetics approach to deciphering the effect of dosage variation on leaf morphology in Populus. Plant Cell 33, 940–960 (2021).

42. N. M. Reid, et al., The genomic landscape of rapid repeated evolutionary adaptation to toxic pollution in wild fish. Science 354, 1305–1308 (2016).

43. P.-I. Zervakis, et al., Genomic studies in Linum shed light on the evolution of the distyly supergene and the molecular basis of convergent floral evolution. New Phytol. 247, 2964–2981 (2025).

44. H. Xue, Y. Gong, S. I. Wright, S. C. H. Barrett, The genomic basis of the tristylous floral polymorphism: Evidence for a role of gene duplications in a region of restricted recombination. Mol. Biol. Evol. 42, msaf170 (2025).

45. D. Charlesworth, The status of supergenes in the 21st century: recombination suppression in Batesian mimicry and sex chromosomes and other complex adaptations. Evol. Appl. 9, 74–90 (2016).

46. N. B. Haghani, et al., Insertion of an invading retrovirus regulates a novel color trait in swordtail fish. [Preprint] (2025). Available at: https://www.biorxiv.org/content/10.1101/2025.11.07.687308v1 [Accessed 19 March 2026].

47. L. Liu, et al., Induced and natural variation of promoter length modulates the photoperiodic response of FLOWERING LOCUS T. Nat. Commun. 5, 4558 (2014).

48. C. Mérot, R. A. Oomen, A. Tigano, M. Wellenreuther, A roadmap for understanding the evolutionary significance of structural genomic variation. Trends Ecol. Evol. 35, 561–572 (2020).

49. M. Göktay, A. Fulgione, A. M. Hancock, A new catalog of structural variants in 1,301 A. thaliana lines from Africa, Eurasia, and North America reveals a signature of balancing selection at defense response genes. Mol. Biol. Evol. 38, 1498–1511 (2021).

50. T. Hämälä, et al., Genomic structural variants constrain and facilitate adaptation in natural populations of Theobroma cacao, the chocolate tree. Proc. Natl. Acad. Sci. 118, e2102914118 (2021).

51. R. Huang, et al., TMK-PIN1 drives a short self-organizing circuit for auxin export and signaling in Arabidopsis. Dev. Cell 61, 73–84.e6 (2026).

52. H. Zheng, et al., Auxin biosynthesis, transport, signaling, and its roles in plant leaf morphogenesis. Plants 15, 72 (2025).

53. Y. Xiong, Y. Jiao, The diverse roles of auxin in regulating leaf development. Plants 8, 243 (2019).

54. T. Huang, et al., Arabidopsis KANADI1 acts as a transcriptional repressor by interacting with a specific cis-element and regulates auxin biosynthesis, transport, and signaling in opposition to HD-ZIPIII Factors. Plant Cell 26, 246–262 (2014).

55. R. A. Kerstetter, K. Bollman, R. A. Taylor, K. Bomblies, R. S. Poethig, KANADI regulates organ polarity in Arabidopsis. Nature 411, 706–709 (2001).

56. R. Solano, A. Stepanova, Q. Chao, J. R. Ecker, Nuclear events in ethylene signaling: a transcriptional cascade mediated by ETHYLENE-INSENSITIVE3 and ETHYLENE-RESPONSE-FACTOR1. Genes Dev. 12, 3703–3714 (1998).

57. O. Lorenzo, R. Piqueras, J. J. Sánchez-Serrano, R. Solano, ETHYLENE RESPONSE FACTOR1 integrates signals from ethylene and jasmonate pathways in plant defense. Plant Cell 15, 165–178 (2003).

58. M.-C. Cheng, P.-M. Liao, W.-W. Kuo, T.-P. Lin, The Arabidopsis ETHYLENE RESPONSE FACTOR1 regulates abiotic stress-responsive gene expression by binding to different cis-acting elements in response to different stress signals. Plant Physiol. 162, 1566–1582 (2013).

59. C. Xu, E. R. Moellering, J. Fan, C. Benning, Mutation of a mitochondrial outer membrane protein affects chloroplast lipid biosynthesis. Plant J. 54, 163–175 (2008).

60. L. Li, et al., Arabidopsis DGD1 SUPPRESSOR1 Is a subunit of the mitochondrial contact site and cristae organizing system and affects mitochondrial biogenesis. Plant Cell 31, 1856–1878 (2019).

61. S. Kimura, et al., CRK2 and C-terminal phosphorylation of NADPH oxidase RBOHD regulate Reactive Oxygen Species production in Arabidopsis. Plant Cell 32, 1063–1080 (2020).

62. G. A. Henry, J. R. Stinchcombe, Predicting fitness-related traits using gene expression and machine learning. Genome Biol. Evol. 17, evae275 (2025).

63. N. C. Durand, et al., Juicer provides a one-click system for analyzing loop-resolution Hi-C experiments. Cell Syst. 3, 95–98 (2016).

64. N. C. Durand, et al., Juicebox provides a visualization system for Hi-C contact maps with unlimited zoom. Cell Syst. 3, 99–101 (2016).

65. E. Linck, C. J. Battey, Minor allele frequency thresholds strongly affect population structure inference with genomic data sets. Mol. Ecol. Resour. 19, 639–647 (2019).

66. P. Danecek, et al., The variant call format and VCFtools. Bioinformatics 27, 2156–2158 (2011).

67. C. C. Chang, et al., Second-generation PLINK: rising to the challenge of larger and richer datasets. GigaScience 4, s13742-015-0047–8 (2015).

68. N. Patterson, A. L. Price, D. Reich, Population structure and eigenanalysis. PLOS Genet. 2, e190 (2006).

69. R. M. Francis, pophelper: an R package and web app to analyse and visualize population structure. Mol. Ecol. Resour. 17, 27–32 (2017).

70. P. Danecek, et al., Twelve years of SAMtools and BCFtools. GigaScience 10, giab008 (2021).

71. B. L. Browning, Y. Zhou, S. R. Browning, A one-penny imputed genome from next-generation reference panels. Am. J. Hum. Genet. 103, 338–348 (2018).

72. R Core Team, R: A language and environment for statistical computing. (2024). Deposited 2024.

73. B. S. Pedersen, A. R. Quinlan, Mosdepth: quick coverage calculation for genomes and exomes. Bioinformatics 34, 867–868 (2018).

74. G. Pertea, M. Pertea, GFF Utilities: GffRead and GffCompare. F1000Research 9, ISCB Comm J-304 (2020).

75. W. Shen, S. Le, Y. Li, F. Hu, SeqKit: A cross-platform and ultrafast toolkit for FASTA/Q file manipulation. PLOS ONE 11, e0163962 (2016).

76. S. F. Altschul, W. Gish, W. Miller, E. W. Myers, D. J. Lipman, Basic local alignment search tool. J. Mol. Biol. 215, 403–410 (1990).

77. J. M. Flynn, et al., RepeatModeler2 for automated genomic discovery of transposable element families. Proc. Natl. Acad. Sci. 117, 9451–9457 (2020).

78. A. F. A. Smit, R. Hubley, P. Green, RepeatMasker Open-4.0. (2013). Deposited 2015 2013.

79. B. Bushnell, Dedupe: A tool for removing duplicate or contained sequences. BBTools. (2023). Deposited 2023.

80. K. J. Hoff, A. Lomsadze, M. Borodovsky, M. Stanke, “Whole-genome annotation with BRAKER” in Gene Prediction: Methods and Protocols, M. Kollmar, Ed. (Springer, 2019), pp. 65–95.

81. J. T. Lovell, et al., GENESPACE tracks regions of interest and gene copy number variation across multiple genomes. eLife 11, e78526 (2022).

82. B. Buchfink, K. Reuter, H.-G. Drost, Sensitive protein alignments at tree-of-life scale using DIAMOND. Nat. Methods 18, 366–368 (2021).

83. D. M. Emms, S. Kelly, OrthoFinder: phylogenetic orthology inference for comparative genomics. Genome Biol. 20, 238 (2019).

84. D. M. Emms, S. Kelly, OrthoFinder: solving fundamental biases in whole genome comparisons dramatically improves orthogroup inference accuracy. Genome Biol. 16, 157 (2015).

85. Y. Wang, et al., MCScanX: a toolkit for detection and evolutionary analysis of gene synteny and collinearity. Nucleic Acids Res. 40, e49 (2012).

86. H. Pagès, P. Aboyoun, R. Gentleman, S. DebRoy, Biostrings: Efficient manipulation of biological strings. (2024). 10.18129/B9.bioc.Biostrings. Deposited 2024.

87. H. Wickham, ggplot2: Elegant Graphics for Data Analysis (Springer-Verlag New York, 2016).

88. SRA Toolkit Development Team, ncbi/sra-tools. (2026). Deposited 8 April 2026.

89. S. Andrews, FastQC: A quality control tool for high throughput sequence data. (2010). Available at: https://www.bioinformatics.babraham.ac.uk/projects/fastqc/ [Accessed 7 November 2025].

90. P. Ewels, M. Magnusson, S. Lundin, M. Käller, MultiQC: summarize analysis results for multiple tools and samples in a single report. Bioinformatics 32, 3047–3048 (2016).

91. F. Krueger, et al., TrimGalore: v0.6.10. (2023). 10.5281/zenodo.7598955. Deposited 2 February 2023.

