## Supplement including: supporting methods, supporting results, Figures S1-S11, Tables S1-S5, Legends for Datasets S1-S5 for "A newly arisen indel governs a leaf shape polymorphism in the Ivy Leaf Morning Glory (*Ipomoea hederacea*)"

**This PDF file includes:**

Supporting Methods
Supporting Results
Figures S1 to S11
Tables S1 to S5
Legends for Datasets S1 to S5
SI References

**Other supporting materials for this manuscript include the following:**

Datasets S1 to S5

**Supporting Information Text**

### Supporting Methods

Seed families included in analyses

After accounting for the individuals we excluded and loss of heterozygosity at the leaf shape locus, we had 123 seed families (61 lobed individuals, 61 entire-shaped individuals, and 1 heterozygote) from a total of 55 populations with 13 entire-shaped populations, 22 lobe-shaped populations, and 20 polymorphic populations (nine polymorphic populations with five individuals, ten polymorphic populations with four individuals, and one polymorphic population with three individuals). We excluded one seed family that had a higher genome-wide levels of heterozygosity than expected for F3 selfed individuals. We calculated heterozygosity with vcftools v0.1.16 (1) and bcftools v1.10.2 (2) using the biallelic SNPs VCF that was generated according to the "Aligning to reference genome and calling SNPs" section below. We further excluded seed families that were noted as homozygotes at the leaf shape locus by (3) but had at least one replicate with leaves of the alternative phenotype.

Genome assembly and annotation:

We chose a lobed seed family, VA12, from a polymorphic population in the middle of the range for the reference genome. Tissue samples for high molecular weight extractions and short read RNA sequencing were collected from a single VA12 individual that was a full sibling of the individuals included in the phenotyping common garden. For RNA extractions, we collected tissue samples from young leaf tissue, old leaf tissue, sepals, the main stem, stem tips, fruit, root, root tips, flower buds. Tissue was placed in liquid nitrogen immediately after collection and placed into a -80°C freezer until extractions. We used QIAGEN RNeasy plant mini kit for RNA extractions. Sequencing libraries were constructed using an Illumina TruSeq Stranded mRNA Library Prep kit and a IDT for Illumina TruSeq UD Indexes Kit using standard protocols. Libraries were sequenced on an Illumina NovaSeq 6000 instrument using paired ends and a read length of 150 basepairs. High Molecular Weight DNA was isolated from flash-frozen leaf tissue using the Circulomics Nanobind kit, sized on an Agilent Femto Pulse, and sheared to a 20 kb mean using a Diagenode MegaRuptor 3. PacBio SMRTbell libraries were prepared and lower-end size selected using a Pippin Prep, and sequenced with 30 hour movies on a PacBio Sequel-II across two flow cells in CCS/HiFi mode. Additionally, a Dovetail Omni-C library was constructed from 1 g of the same flash-frozen leaf tissue according to manufacturers instructions, and sequenced on an Illumina NovaSeq X with PE150 reads. HiFi reads were assembled in hifiasm v.0.19.3-r572 (4–6) with Omni-C integration. Given the high homozygosity of the line, only a single haplotype was assembled and scaffolded using YaHS v.1.2.2 (7). Omni-C reads were aligned to hifiasm contigs using bwa mem with flags "-5SP", contigs were manually ordered and re-oriented based on the contact map, and chromosomes were renamed against the *Ipomoea nil* genome in juicer_tools v1.22.01 (8) and juicebox v2.20.00 (9). The *I. hederacea* genome was repeat-masked using RepeatModeler v2.0.3 (10) and RepeatMasker v4.1.4 (11). RNA-seq short reads were cleaned with trimmomatic v.039 (12), deduplicated with nubeam-dedup v25dd385 (13), and assembled with IDBA-tran v1.1.1 (14). The RNA-seq contigs and OrthoDB v. 11 Viridiplatae proteins downloaded from https://bioinf.uni-greifswald.de were used as evidences in the genome annotation with BREAKER genome annotator (15). We ran BUSCO v6.0.0 (16) against OrthoDB v. 12 Viridiplatae to assess annotation completeness.

DNA short-read whole genome extractions, library prep, and sequencing:

We conducted whole-genome short-read resequencing for the 123 seed families. We grew up a single individual from each seed family that was previously selfed for at least three generations in a common greenhouse environment to collect tissue for DNA extraction. Young leaf tissue was harvested from each individual, immediately placed into liquid nitrogen, and then stored in a -80°C freezer until DNA extractions. We used QIAGEN DNeasy plant mini kit to perform DNA extractions. Illumina sequencing libraries were prepared from 1.5 μg of DNA sheared to 350 bp using a Covaris LE220 instrument. DNA fragments were bead cleaned, end repaired, and size selected to remove large and small fragments. After adenylation, adaptors were ligated according to the Illumina TruSeq PCR-Free DNA Library Prep Kit Protocol. Libraries were sequenced on an Illumina NovaSeq 6000 instrument using paired ends and a read length of 150 bp.

Aligning to reference genome and calling SNPs:

We used fastqc v0.12.0 (17) and multiqc v1.19 (18) to assess short read sequencing quality that indicated there were some adapter sequences, a skewed representation of base pairs on the 3' end, and an overrepresentation of poly-G sequences. We trimmed adapter sequences using the *--illumina* flag and then hard clipped 6 based pairs on the 3’ end of both forward and reverse strands using TrimGalor v0.6.10 (19). The reference genome was indexed using bwa index and we aligned the short read sequencing to the reference genome using bwa mem v0.7.17 (20). We used samtools view to convert aligned .sam files to .bam files; samtools fixmate to add mate score tags; samtools sort to sort bam files be coordinates; samtools markdup to mark duplicated reads; and then finally samtools index to index all bam files v1.18 (2). We then used qualimap v2.3 (21), multiqc, and samtools flagstat to assess read mapping quality. To generate a VCF file, we used bcftools mpileup v.1.19 (2) with probabilistic realignment disabled for the computation of base alignment quality and we added information tags for allelic depth and number of high-quality bases. We called variants using bcftools call for all chromosomes separately and then we combined VCFs using bcftools concat. We used bcftools view to remove seed families excluded from analyses and updated all info tags using the bcftools plugin “+fill-tags”. We then used bcftools filter to remove all non-variant sites, sites with a phred quality score <30, sites with a mapping quality <30, sites missing calls for >10% of all seed families, and sites with a minimum mean depth below 10x and above 40x. We then used bcftools view to remove all genetic variants that were not biallelic SNPs. The VCF used in downstream analyses, unless otherwise stated, had a total of 2,990,321 biallelic SNPs after filtering.

Common Garden Phenotyping:

Seeds used in the common garden experiment were F3 individuals produced from the same selfed parents that were used for the collecting sequencing for tissue extractions. Each seed family had one replicate per block across five blocks. Seeds were scarified to sync germination and then planted in conetainers with promix soil. The greenhouse temperature was 27-29°C with 16-hour day lengths to facilitate vegetative growth throughout the experiment. We misted plants daily for the first nine days and then bottom watered every other day for the next two weeks. From day 23 onwards greenhouse benches were continually flooded with water. Biocontrols (*Stratiolaelaps scimitus*, *Steinernema feltiae*, *Neoseiulus cucumeris*) for pest management and 20-20-20 Miracle-Gro fertilizer were applied as needed throughout the experiment.

Leaf area, circularity, and roundness: The sixth true leaf from each individual was used to measure leaf shape. If the sixth leaf showed any signs of damage or yellowing, then the closest healthy leaf (i.e., the 5th or 7th leaf) was used instead. A photo was taken of the leaves laid flat on a 1 cm x 1 cm grid background. ImageJ was used to capture leaf outlines and calculate leaf area, circularity, and roundness.

Leading vine length: We measured the length of the leading vine on days 15 for all five blocks and day 22 for blocks one and two. We removed stem length measurements for three individuals (DE240_2, MD202_1, and TN669_1) that had smaller length measurements for day 22 than day 15.

Leaf counts: We performed a total of four leaf counts spaced one week apart on days 14, 21, 28 and 35 of the experiment. We decided to not exclude data for individuals with higher values for previous leaf counts because it is possible that the loss of leaves between counts could cause a lower number of leaves at a later time point.

Trichome density: The tenth true leaf from each individual was used to measure trichome density. If the tenth leaf showed any signs of damage or yellowing, then the closest healthy leaf was used instead. We took a single 0.6 mm diameter sample from the top right of each leaf using a hole punch. Leaf cores were imaged under a dissecting microscope. We then used the ImageJ counter function to count the trichomes for each sample.

Relative water content: To calculate relative water content we used the equation $RWC = \frac{(W-DW)}{(TW-D\ \ )}$ where W is fresh weight, TW is turgid weight, and DW is dry weight. A fully expanded leaf was taken from the top of each individual and immediately weighed for fresh weight measurement. Leaves were then left to soak in water for 4 hours. After which the leaves were removed from the water, patted dry with a paper towel, and then weighed for turgid weight measurement. Leaves were then placed in a drying oven at 60°C for 72 hours and then weighed once more to determine leaf dry weight. We staggered measurements across multiple days by only measuring one block a day (eg. Day 1: W and TW for Block 1, Day 2: W and TW for Block 2… Day 4: W and TW for Block 4 and DW for Block 1).

Leaf surface temperature and stomatal conductance: We used a porometer (Decagon Devices, SC-1 Model) to measure the surface temperature and stomatal conductance of the upper leaf surface. Given that leaf surface temperature and stomatal conductance can vary throughout the day, we collected measurements from 12pm to 4pm when stomatal conductance should be at its highest. Data was collected across four days with half a block being measured per day.

To determine if lobing is associated with the other phenotypic traits we measured, we used two-sided t-tests and Bonferroni corrected p-values to account for multiple comparison testing ($p_{adj} = p_{raw}$*12). We used seed family means for each seed family for the t-test and we excluded the heterozygote seed family (TN682) because the phenotypes of a heterozygote seed family will include a mixture of all three possible leaf shape phenotypes.

<u>Calculating LD decay:</u>
We used PLINK v1.9b_6.21-x86_64 (22) to calculate the squared correlation coefficient ($r^2$) for 10% of randomly selected loci. We restricted the pairwise $r^2$ calculations to SNPs that were no more than 1000 kb apart and had no more than 100 variants between them. We then calculated the average $r^2$ for bins of 1000 bp to generate an LD decay plot (Figure S1). Based on the LD decay plot, we then chose to LD prune the biallelic VCF to remove any SNPs within a 250 kb window with an $r^2 > 0.1$ for the population genomic analysis. We had a total of 137,699 SNPs remaining after filtering for singletons, non-neutral sites, and LD.

<u>Aligning to reference genome without indel:</u>
We aligned individuals to a reference of Chromosome 2 with the indel removed to determine if the structural variant is a simple deletion in entire-shaped individuals or possibly a more complex structural variant. We first determined the putative indel breakpoints by visually inspecting a subset of entire-shaped and lobed individuals in IGV. We used samtools faidx to create two subset reference genomes: one of Chromosome 2 before the indel (Chr2:1-3625421) and another of Chromosome 2 after the indel (Chr2:3742102-43274491). We then concatenated the fasta files representing the sequence before and after the indel to create a new pseudo-reference genome of Chromosome 2 that is missing the indel. We aligned entire-shaped and lobed individuals to the pseudo-reference genome as described in the “Aligning to reference genome and calling SNPs” methods section. We used bedtools intersect v2.31.0 (23) and samtools view v1.18 to count the reads that mapped across the entire 10 bp region before the removed indel (Chr2:3625410-3625420), overlapping with the removed indel region (Chr2:3625416-3625426), and after the removed indel (Chr2:3625423-3625433). We conducted a t-test comparing read counts between entire-shaped and lobed individuals for the 10 bp before, overlapping, and after the removed indel region in R.

Visualisations:
Visualisations were generated in R v4.4.0 (24) using R studio v2024.9.999.999 (25) using R packages ggplot2 v4.0.0 (26), ggpubr v0.6.2 (27), gggenes v0.5.1, patchwork v1.3.2 (28), ggtext v0.1.2 (29), ggfittext v0.10.2 (30), and gridExtra v2.3 (31) unless otherwise stated.

### Supporting Results

Leaf shape GWAS hits other than Chromosome 2
Along with the highly significant GWAS peaks on Chromosome 2 for leaf shape, we also found barely significant peaks on Chromosomes 6, 7, and 14 across the various GWAS analyses we performed. In the GWAS for lobed vs entire shapes as a binary phenotype (n=122), there were GWAS hits on Chromosome 2 and 7 (Figure 2). The GWAS hit on Chromosome 7 was much less significant than the hit on Chromosome 2 but it still surpassed the Bonferroni significance threshold (Figure 2A). The significant region only spans 145 bp (Chr7:11966510-11966655) with only four SNPs that are above the Bonferroni significance threshold. All four SNPs have low allele frequencies ranging from 0.098-0.115. For the leaf circularity GWAS that included all individuals (n=122), there are GWAS hits on Chromosome 2, 6, and 7 (Supplementary Figure S7A-B). The peak on Chromosome 7 has only four SNPs within a 145 bp region (Chr7: 11966510 - 11966655) with low minor allele frequencies that ranged from 0.098-0.115). There was only a single SNP on Chromosome 6 (Chr6: 7284604) that was significantly associated with leaf circularity and also had a low minor allele frequency of 0.053. In the leaf circularity GWAS that only included lobed individuals (n=61), there are GWAS hits on Chromosome 2, 7, and 14 (Figure S7C-D). Only two SNPs that were 65 bp apart passed the FDR 0.05 significance threshold on Chromosome 7 (Chr7:11966510-11966575). Both of the significant SNPs on Chromosome 7 were also significant in the lobed vs entire binary GWAS. There is only one SNP on Chromosome 14 (Chr14: 41393623) that surpassed the 0.05 FDR significance threshold. The low allele frequencies and relatively lower peaks compared to Chromosome 2 suggest that the barely significant peaks on Chromosomes 6, 7, and 14 are likely spurious correlations.

Adaptive latitudinal clines in *I. hederacea*
Overall, we find a lack of neutral population genetic structure in *I. hederacea* suggesting that the observed latitudinal clines in the species are likely due to selection. The best fit ADMIXTURE model does not differentiate any of the samples into distinct clusters. Additionally, we found very weak clustering in PC space for the principal components that explained the most amount of neutral genetic variation (Figure 1C-G). A previous study using 173 AFLP loci also found little evidence of population structure (3). However, a follow-up study using sanger sequencing of seven nuclear genes did find evidence of isolation-by-distance in *I. hederacea* but neutral genetic variation was not geographically structured (32). Together the results support previous hypotheses that long distance human-mediated dispersal combined with high rates of population turnover has resulted in weak geographic population structure (3, 32) similar to other *Ipomoea* species (33, 34). Previous studies have reported latitudinal clines in flowering time, floral morphology, biomass, and growth rate in addition to leaf shape in *I. hederacea* (35, 36). The lack of geographic population structure reported in multiple studies, using multiple genetic markers and technologies, suggests that the latitudinal phenotypic clines in *I. hederacea* are due to selection. However, it is worth noting that previous studies looking at population structure in *I. hederacea* (3, 32) are not completely independent of this study as we used an overlapping set of seed families with more up-to-date sequencing technology and better representation of genome-wide genetic variation. The potential mechanisms of selection on leaf shape, particularly forces that favor lobed genotypes in the northern part of the eastern North American range, nonetheless remain elusive. For example, in a series of experiments in North and South Carolina, in the polymorphic portion of *I. hederacea*'s range, Bright and Rausher (2008) found balancing selection in the form of heterozygote advantage for fitness at two sites, and selection favoring the lobed genotype in 1 site-year combination. Their experiments, however, were unable to

characterize the ecological mechanisms leading to fitness advantages of the lobed genotypes in that year. A handful of studies in the northern part of the eastern North American range and beyond, have detected strong selection in the field on herbivore resistance, size, growth rate, and flowering phenology, that did not differ between leaf shape genotypes (37, 38). Conversely, several studies have detected differences a host of physiological and biotic interaction traits [night-time thermoregulation (39), stomatal conductance in the field on sunny days (40), herbivory preferences (41), pathogen resistance (42), and leaf microbiome assembly (43)] that did not lead to differences in survival or reproduction. Our finding that the indel contains several genes that the cline may be due to the indel's effects on multiple ecologically important traits (either simultaneously, or in combination, depending on how selection acts in any given site or year), rather than having a single ecological cause.

**Figures**

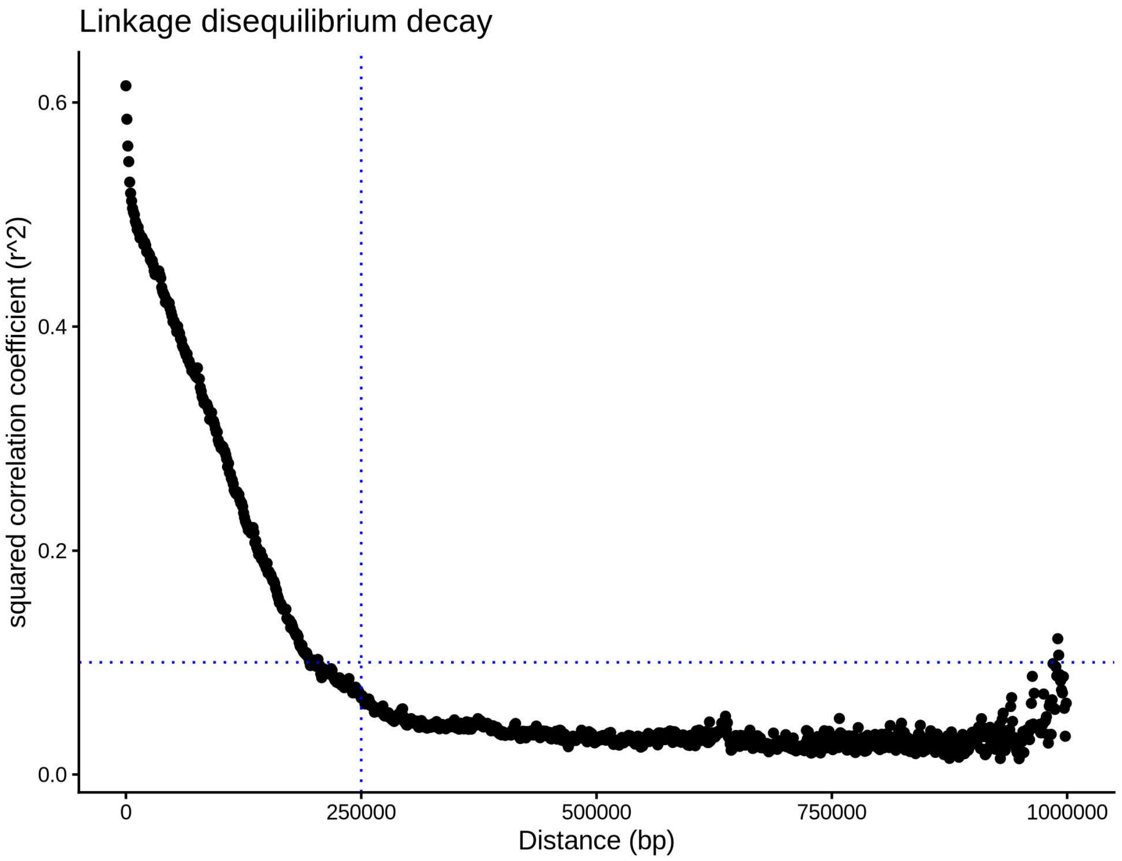


**Figure S1.** Scatter plot of the mean squared correlation coefficient for 1 kb bins. The blue dotted lines represent the LD pruning thresholds of 250 kb (vertical line) and $r^2 = 0.1$ (horizontal line).

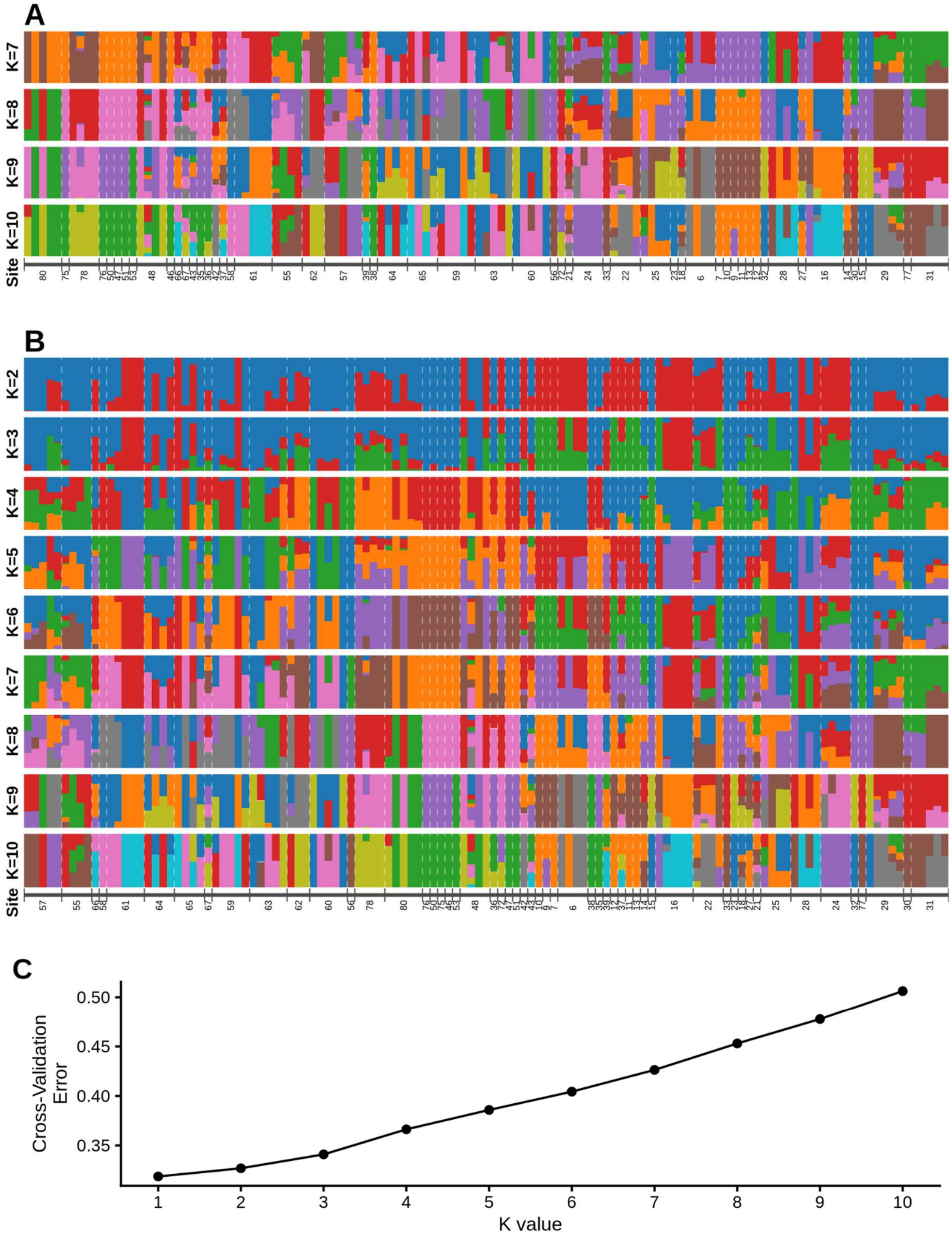


**Figure S2. A)** Neutral genetic diversity ADMIXTURE plots of K7-10 with populations ordered by latitude and **B)** with populations ordered by longitude for K2-10. **A-B)** Individuals from different populations are separated by white dashed lines and the population sampling sites are denoted underneath the structure plots. **C)** ADMIXTURE cross validation error for K values 1-10.

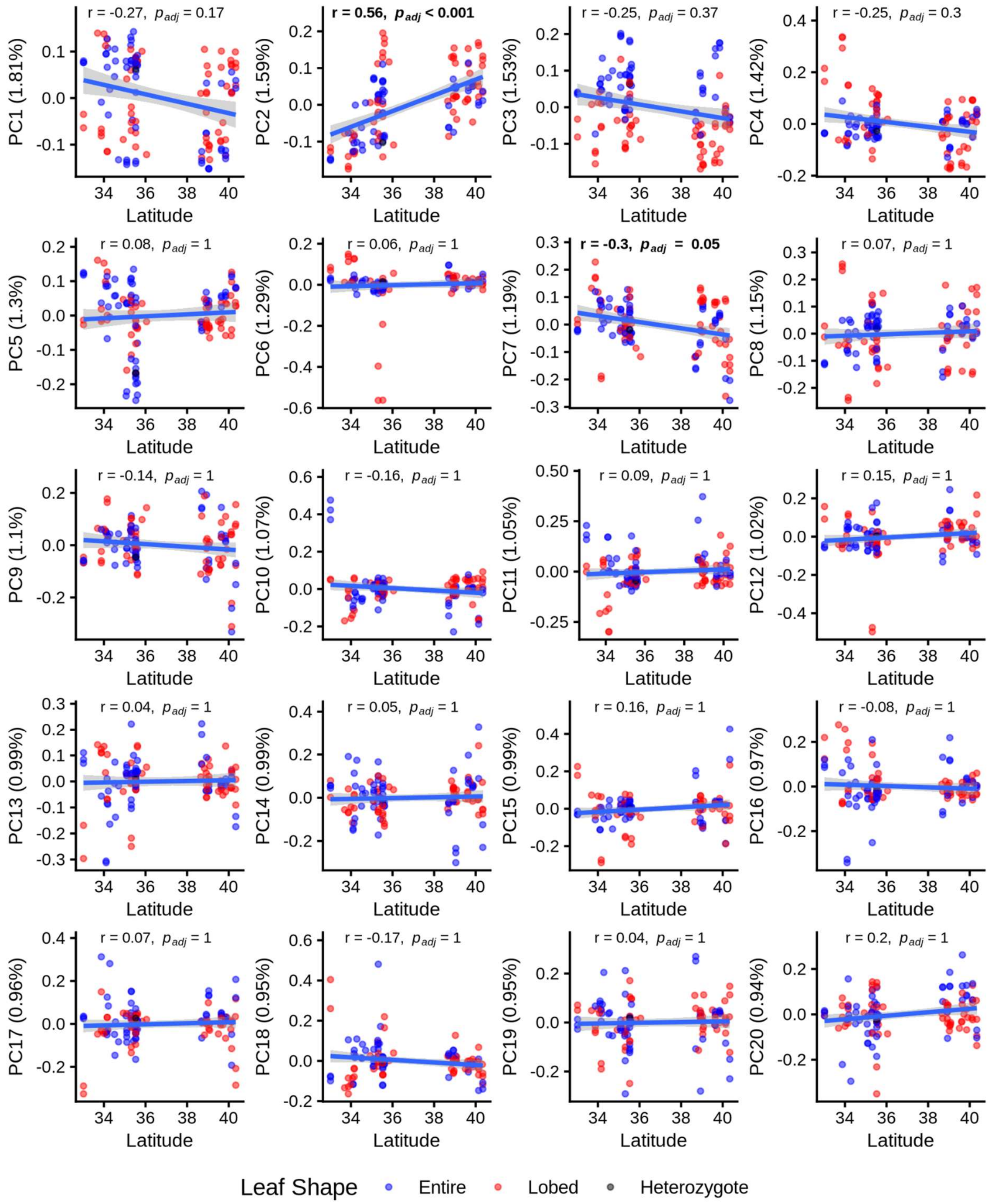


**Figure S3.** Principal components 1-20 plotted against latitude. Each data point represents a seed family and the color of the point indicates the seed family leaf shape. Pearson correlation coefficients and Bonferroni corrected p-values ($p_{adj} = p_{raw}$*60) are shown in the plots and statistically significant correlations are bolded. For each plot the blue line indicates the fitted linear model with the 95% confidence interval shown in grey.

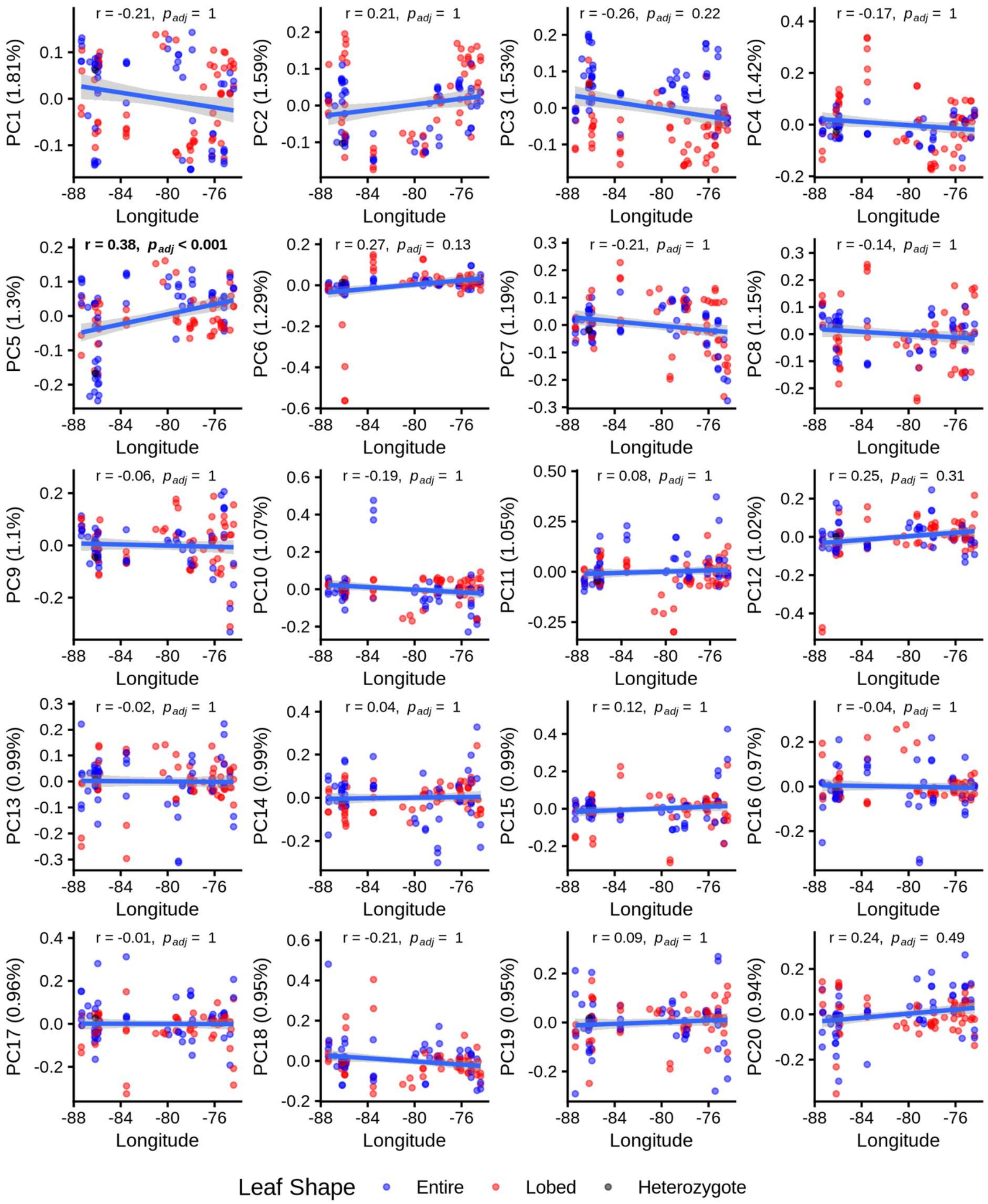


**Figure S4.** Principal components 1-20 plotted against longitude. Each data point represents a seed family and the color of the point indicates the seed family leaf shape. Pearson correlation coefficients and Bonferroni corrected p-values ($p_{adj} = p_{raw}$*60) are shown in the plots and statistically significant correlations are bolded. For each plot the blue line indicates the fitted linear model with the 95% confidence interval shown in grey.

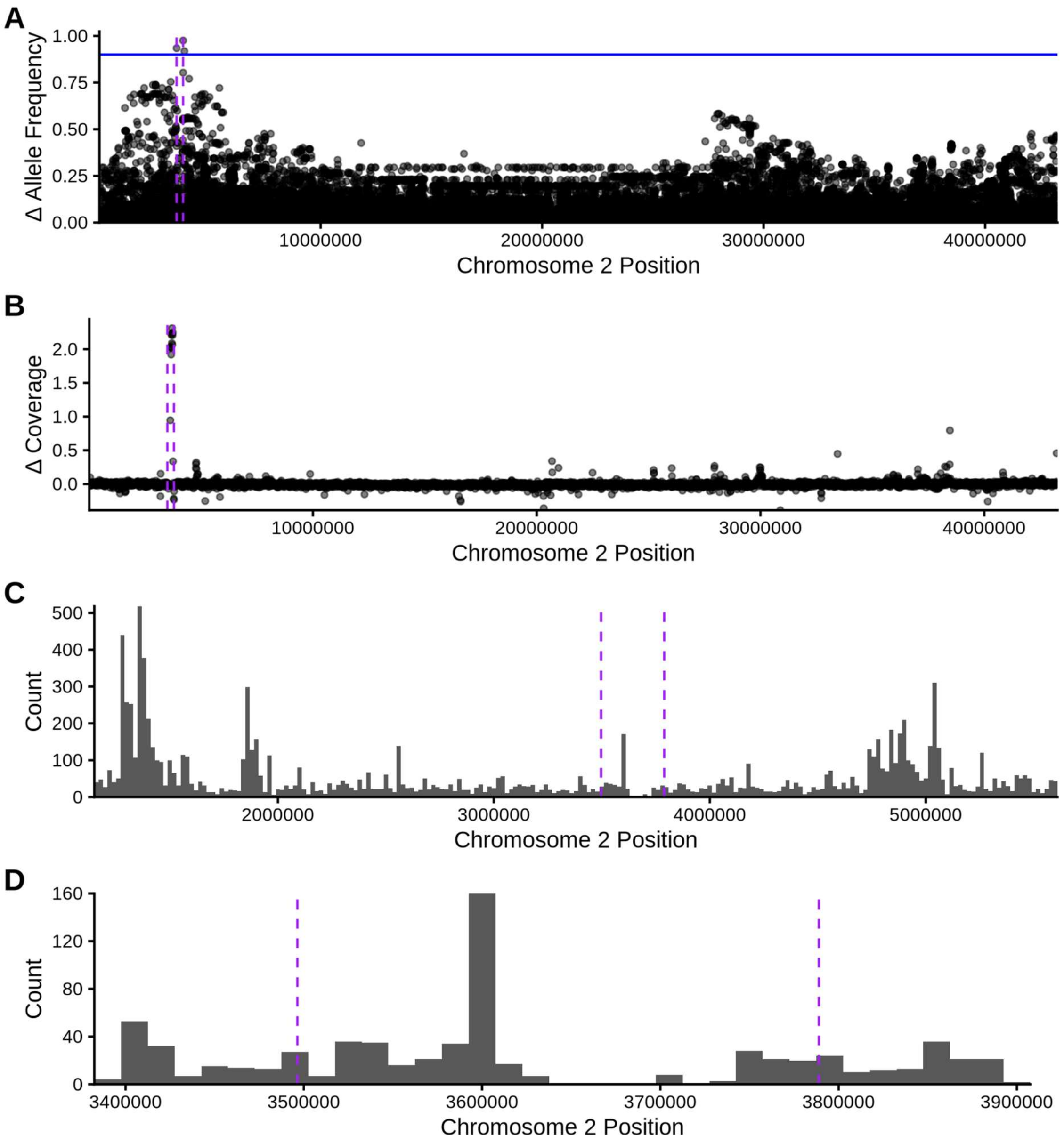


**Figure S5. A)** Manhattan plot of allele frequency differences between leaf shapes across all of Chromosome 2. Blue horizontal line shows the allele frequency difference of 0.9. **B)** Manhattan plot of difference in mean normalized coverage between leaf shapes (lobed - entire) across all of Chromosome 2. **C)** Histogram of SNP density for leaf shape GWAS peak on Chromosome 2 (binwidth = 20 kb). **D)** Zoomed in histogram of SNP density (binwidth = 15 kb) within the region surrounding leading GWAS SNPs. **A-D)** Purple dashed lines mark the locations of the two GWAS SNPs that are most significantly associated with leaf shape.

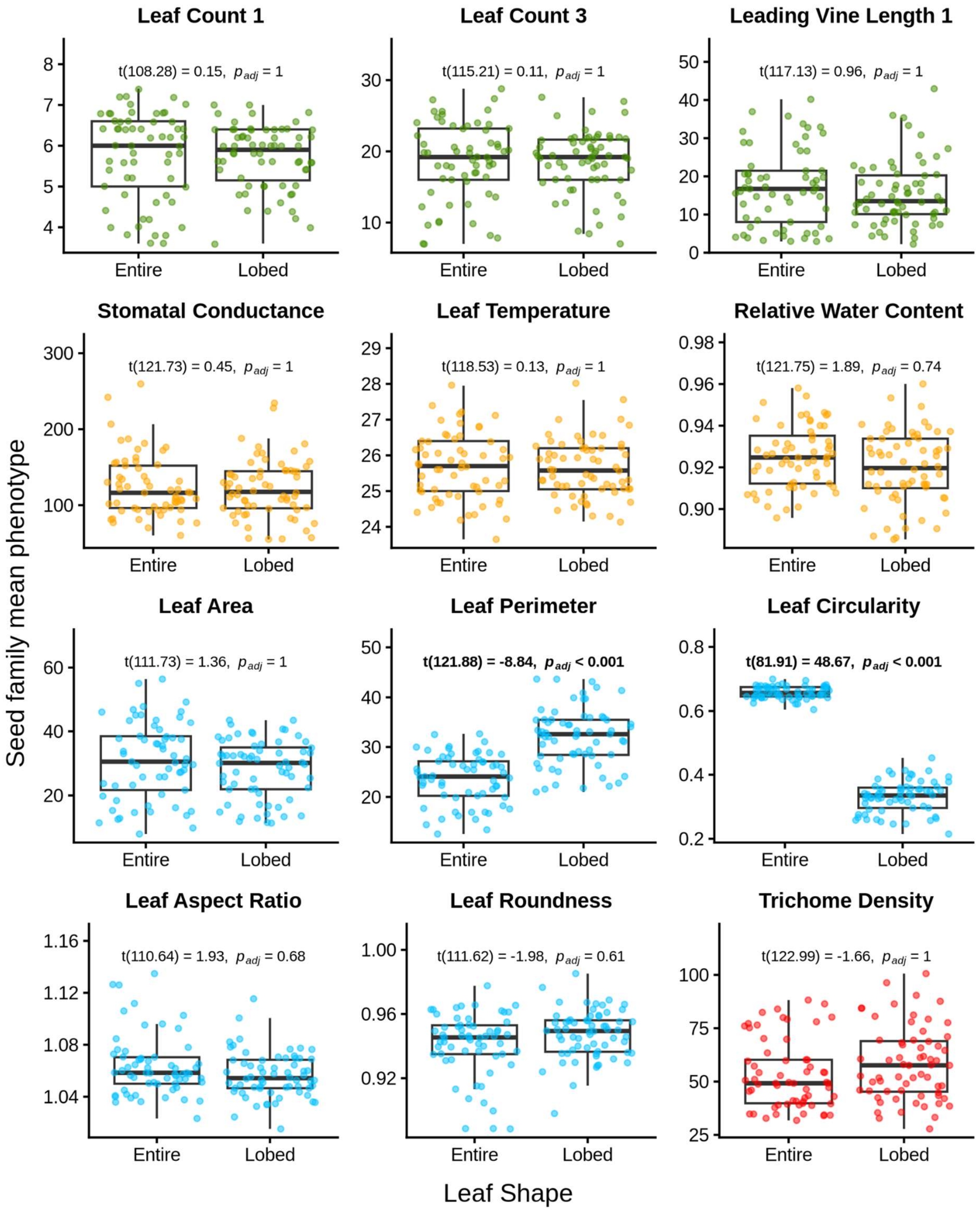


**Figure S6.** Boxplots of differences in proxies for early growth rate (green), ecophysiological (orange), leaf shape (blue), and trichome density (red) traits between lobed and entire-shaped seed families. Each data point represents seed family mean. Two-sided t-test statistics are shown at the top of each box plot including the degrees of freedom, test statistic, and Bonferroni corrected p-value ($p_{adj} = p_{raw}*12$). Statistics representing significant differences between leaf shapes are bolded.

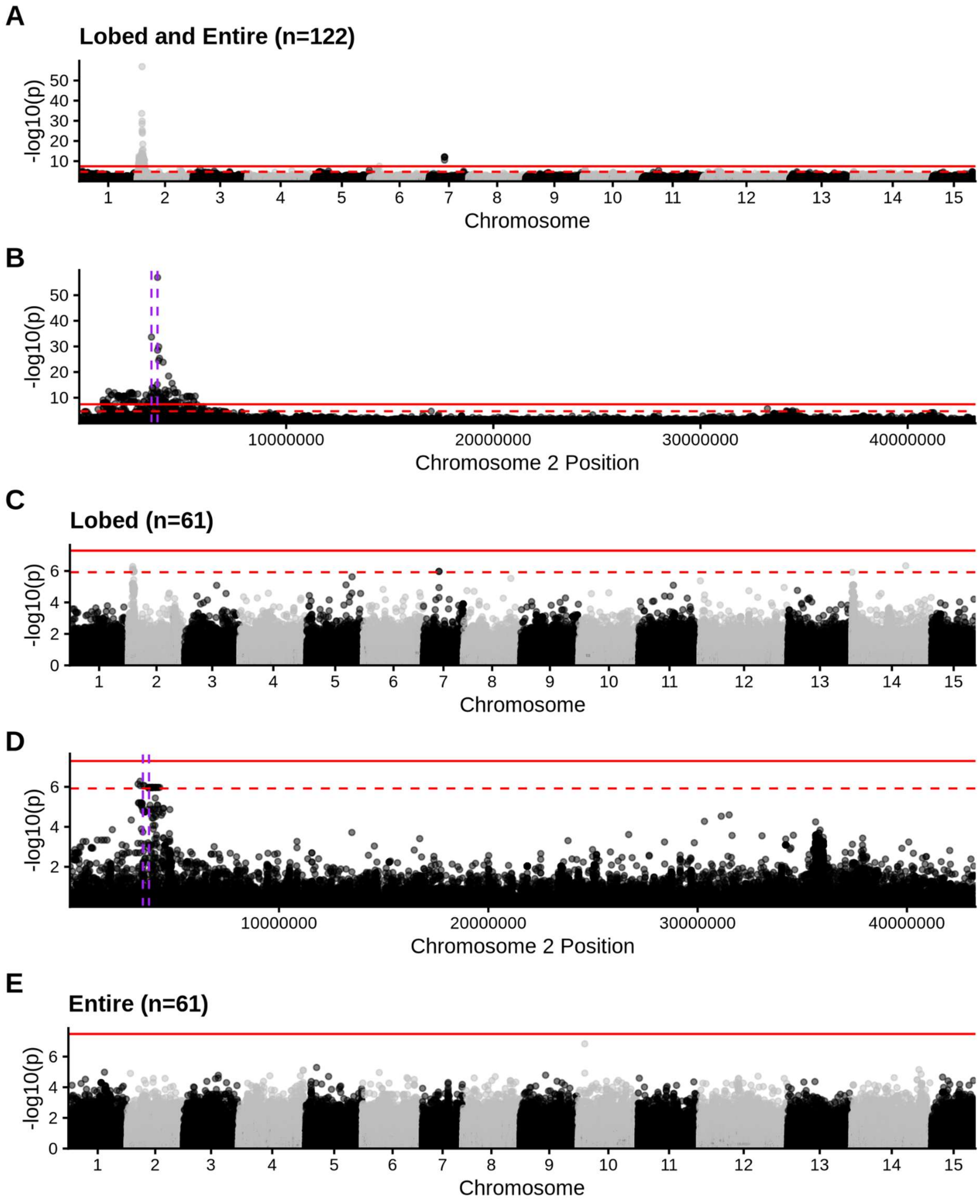


**Figure S7.** Manhattan plots for circularity GWAS including **A-B)** both lobed and entire-shaped individuals (n=122), **C-D)** only lobed individuals (n=61), and **E)** only entire-shaped individuals (n=61). **A, C, E)** Genome-wide manhattan plots. **B, D)** Manhattan plot of Chromosome 2. The purple vertical dashed lines mark the locations of the two GWAS SNPs that are most significantly associated with leaf lobing (as a binary trait) in Figure 3. **A-E)** The red horizontal lines represent the bonferroni corrected (solid) and 0.05 FDR (dashed) significant thresholds. Manhattan plots that had no SNPs with an FDR adjusted p-value < 0.05 do not have an FDR significance threshold plotted.

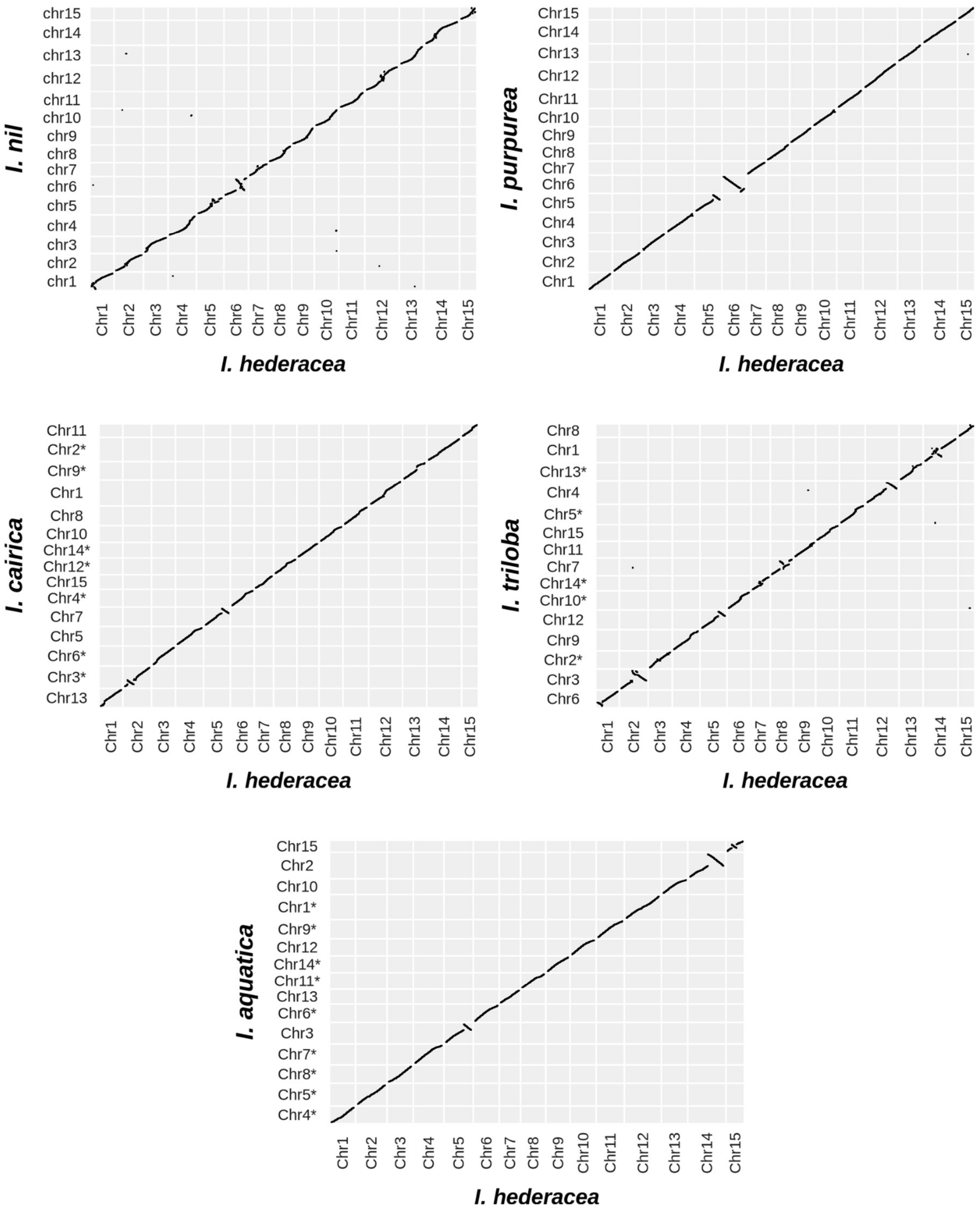


**Figure S8.** Genome-wide dot plots of syntenic orthologs between *I. hederacea* and other *Ipomoea* species plotted by gene order. We excluded smaller contigs from the plots that were not included in the main 15 chromosomes for each species. The order of chromosomes for *I. triloba, I. cairica,* and *I. aquatica* was rearranged to match the chromosome naming order *I. hederacea*. All chromosomes that have been inverted to match the orientation of the *I. hederacea* assembly are marked with an asterisk in the y-axis.

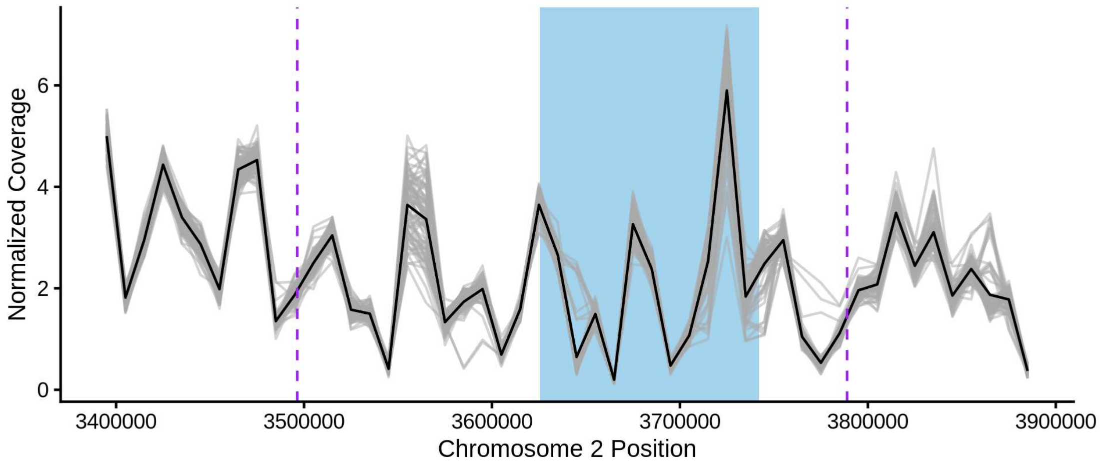


**Figure S9.** Normalized coverage of 78 *I. purpurea* individuals aligned to the *I. hederacea* genome across the leaf shape indel region in *I. hederacea*. Each grey line represents the normalized coverage for an *I. purpurea* individual that was previously sequenced by Gupta et al. 2023 (44). The black line represents the mean normalized coverage for all *I. purpurea* individuals. The blue shaded box represents the indel co-ordinates in *I. hederacea*. Purple dashed lines are the loci of the two SNPs with the highest significance in the *I. hederacea* leaf shape shown in Figure 2.

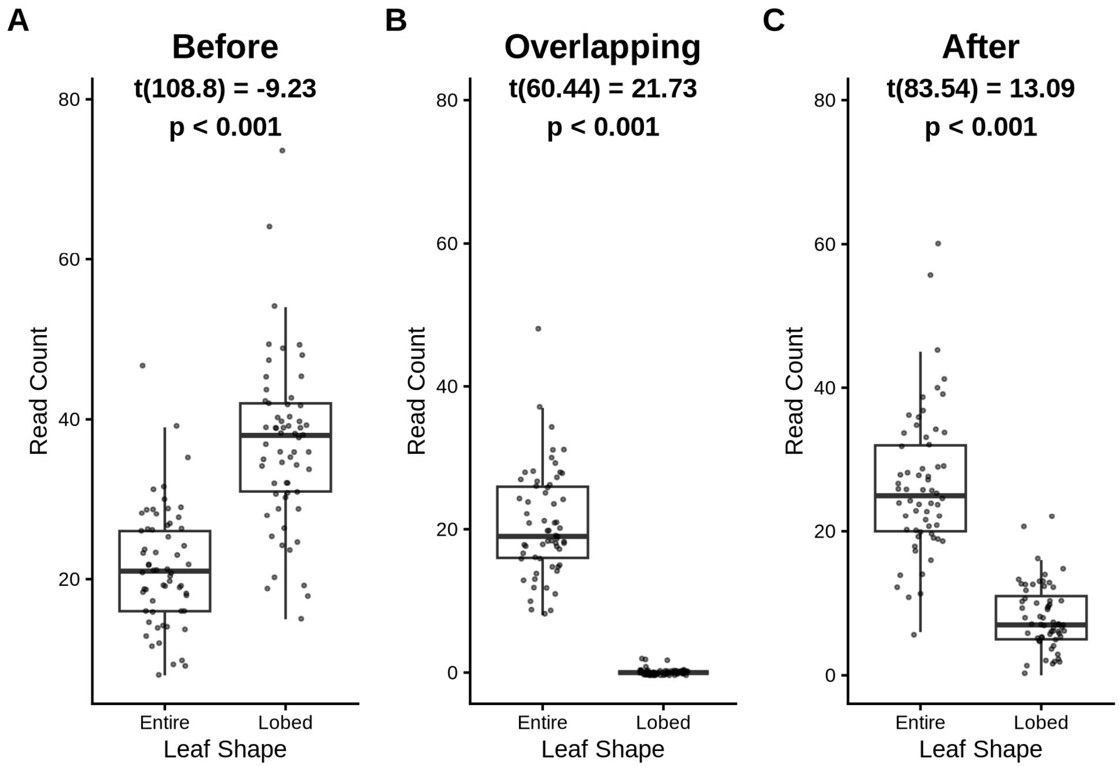


**Figure S10.** Boxplots comparing the number of reads in lobed and entire-shaped individuals that are aligned to a pseudo-reference genome of Chromosome 2 with the indel removed. Read counts include any reads that map continuously across the 10 bp region **A)** before, **B)** overlapping, and **C)** after the removed indel region. Two-sided t-test statistics (degrees of freedom, test statistic, and p-value) are shown at the top of each box plot with significant results in bold.

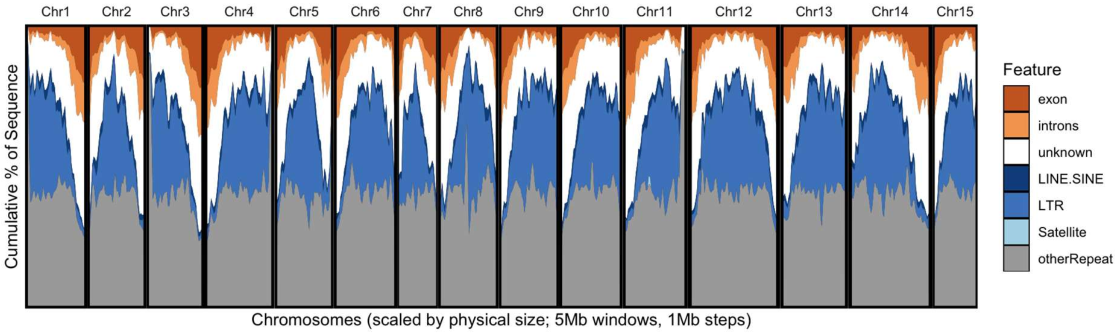


**Figure S11.** Cumulative percentage of sequence for genomic features across all 15 chromosomes for the *I. hederacea* genome annotation and repeat content.

**Tables**

| Species | Complete | Single | Duplicated | Fragmented | Missing |
|---|---|---|---|---|---|
| *Ipomoea hederacea* | 99.6 | 89.7 | 10.0 | 0.0 | 0.4 |
| *Ipomoea nil* | 99.9 | 89.8 | 10.1 | 0.0 | 0.1 |
| *Ipomoea purpurea* | 94.3 | 85.4 | 8.9 | 1.9 | 3.8 |
| *Ipomoea acquatica* | 97.7 | 89.3 | 8.4 | 0.5 | 1.8 |
| *Ipomoea cairica* | 96 | 86.9 | 9.1 | 3.2 | 0.9 |
| *Ipomoea triloba* | 96.1 | 87.5 | 8.6 | 1.5 | 2.4 |

**Table S1.** Busco scores for *Ipomoea* species annotations

**Table S2.** Top BLAST hits for *I. hederacea* genes on the putative leaf shape indel when BLASTed against *A. thaliana* TAIR10 and also the NCBI database.

| ***I. hederacea* gene** | ***A.thaliana* top BLAST hit** | | **NCBI top BLAST hit** | | |
|---|---|---|---|---|---|
| | **Gene** | **e-value** | **Gene** | **Species** | **e-value** |
| g17358 | ethylene response factor 1 | 5.05e-43 | PREDICTED: ethylene-responsive transcription factor 1B-like | *Ipomoea nil* | 2.39e-99 |
| g17359 | dgd1 suppressor 1 | 0.15 | PREDICTED: uncharacterized protein LOC109157732 isoform X2 | *Ipomoea nil* | 4.84e-41 |
| g17360 | ethylene response factor 1 | 2.69e-42 | PREDICTED: ethylene-responsive transcription factor 1B-like | *Ipomoea nil* | 1.20e-62 |
| g17361 | zinc finger protein 4 | 7.03e-23 | zinc finger protein 4-like | *Ipomoea trifida* | 1.63e-69 |
| g17362 | cysteine-rich RLK (RECEPTOR-like protein kinase) 2 | 3.63e-125 | PREDICTED: cysteine-rich receptor-like protein kinase 2 | *Ipomoea nil* | 0.0 |
| g17363 | cysteine-rich RLK (RECEPTOR-like protein kinase) 2 | 4.91e-116 | PREDICTED: cysteine-rich receptor-like protein kinase 2 | *Ipomoea nil* | 0.0 |
| g17364 | Transmembrane amino acid transporter family protein | 1.94e-19 | vacuolar amino acid transporter 1-like isoform X1 | *Ipomoea batatas* | 8.59e-30 |

**Table S3.** Candidate leaf shape genes found within 250 kb upstream or downstream of the two most significant GWAS SNPs.

| ***I. hederacea* gene** | ***A. thaliana* top BLAST hit** | | **NCBI top BLAST hit** | | |
|---|---|---|---|---|---|
| | **Gene** | **e-value** | **Gene** | **Species** | **e-value** |
| g17353 | transmembrane kinase 1 | 0.0 | PREDICTED: receptor protein kinase TMK1-like | *Ipomoea nil* | 0.0 |
| g17384 | Homeodomain-like superfamily protein | 2.63e-71 | PREDICTED: transcription repressor KAN1 isoform X1 | *Ipomoea nil* | 0.0 |

**Table S4.** Welch two sample t-test statistics for PC1-20 and leaf shape (n=122).

| PC | t-statistic | Degrees of freedom | Raw p-value | Bonferroni corrected p-value |
|---|---|---|---|---|
| PC1 | 0.51 | 117.81 | 0.61 | 1 |
| PC2 | -2.27 | 109.97 | 0.03 | 1 |
| PC3 | 9.89 | 119.58 | 3.27e-17 | 1.96e-15 |
| PC4 | -1.50 | 75.33 | 0.14 | 1 |
| PC5 | 0.08 | 95.25 | 0.93 | 1 |
| PC6 | 0.76 | 65.27 | 0.45 | 1 |
| PC7 | -0.49 | 113.93 | 0.63 | 1 |
| PC8 | 0.22 | 98.24 | 0.83 | 1 |
| PC9 | -0.29 | 119.59 | 0.77 | 1 |
| PC10 | -0.11 | 92.44 | 0.91 | 1 |
| PC11 | 2.61 | 119.85 | 0.01 | 0.61 |
| PC12 | 0.34 | 100.78 | 0.73 | 1 |
| PC13 | -0.16 | 118.70 | 0.87 | 1 |
| PC14 | 0.18 | 97.24 | 0.86 | 1 |
| PC15 | 0.65 | 119.77 | 0.51 | 1 |
| PC16 | -1.63 | 115.45 | 0.11 | 1 |
| PC17 | 2.32 | 119.12 | 0.02 | 1 |
| PC18 | 0.83 | 119.99 | 0.41 | 1 |
| PC19 | -0.55 | 111.75 | 0.58 | 1 |
| PC20 | 0.78 | 116.12 | 0.44 | 1 |

**Table S5.** Genomic locations of top BLAST hits from BLASTing *I. hederacea* indel genes to other *Ipomoea* protein databases. Green represents hits found in the same syntenic region, orange indicates hits that are still in the same syntenic region but are the top hits for two *I. hederacea* indel genes, red denotes non-syntenic top BLAST hits.

| ***I. hederacea* Gene ID (Top *A. thaliana* BLAST hit)** | ***I. hederacea*** | ***I. nil*** | ***I. purpurea*** | ***I. triloba*** | ***I. acquatica*** | ***I. cairica*** |
|---|---|---|---|---|---|---|
| g17358.t1 (ethylene response factor 1) | Chr2: 3633990 3634427 | Chr2: 3664062 3664499 | Chr2: 2967079 2967450 | Chr3: 3105516 3105947 | Chr5: 32104101 32104562 | Chr3: 47004889 47005341 |
| g17359.t1 (dgd1 suppressor 1) | Chr2: 3643283 3643864 | Chr5: 16652337 16653102 | Chr2: 18407924 18408558 | Chr1: 17564748 17565941 | Chr13: 17068051 17068623 | Chr6: 22183225 22183838 |
| g17360.t1 (ethylene response factor 1) | Chr2: 3653948 3654400 | Chr2: 36 83940 3684383 | Chr2: 298 2248 2982700 | Chr3: 3105516 3105947 | Chr5: 32104101 32104562 | Chr3: 47004889 47005341 |
| g17361.t1 (zinc finger protein 4) | Chr2: 3671574 3672173 | Chr2: 36 89755 3690456 | Chr2: 298 8294 2989013 | Chr3: 3146487 3147023 | Chr5: 32081354 32082028 | Chr3: 47084415 47085071 |
| g17361.t2 (zinc finger protein 4) | Chr2: 3671460 3672173 | Chr2: 36 89755 3690456 | Chr2: 298 8294 2989013 | Chr3: 3146487 3147023 | Chr5: 32081354 32082028 | Chr3: 47084415 47085071 |
| g17362.t1 (cysteine-rich RLK 2) | Chr2: 3683400 3687093 | Chr2: 37 05063 3708782 | Chr5: 764 6957 7649487 | Chr3: 3154709 3158140 | Chr5: 32074697 32078316 | Chr3: 47104368 47108610 |
| g17363.t1 (cysteine-rich RLK 2) | Chr2: 3715673 3720270 | Chr2: 38 20842 3824919 | Chr2: 305 3706 3058850 | Chr3: 3162111 3168719 | Chr5: 32052731 32056241 | Chr3: 47162957 47168318 |
| g17364.t1 (Transmembrane amino acid transporter family protein) | Chr2: 3731421 3732184 | Chr4: 82 22092 8222763 | Chr4: 639 2700 6393332 | Chr9: 7764364 7764963 | Chr7: 23392963 23393576 | Chr5: 10826070 10826738 |

**Dataset S1 (supplementary_tracy_widom_df.csv).** Tracy-Widom statistic for population genetics PCA. Columns description:

1) eigenvals = Eigen values for each principal component
2) variation_explained = The amount of variation explained by each principal component
3) variation_percentage = The amount of variation explained by each principal component as a percentage
4) tw_significant = If the PC explains a “Significant” or “Not Significant” amount of variation based on the tracy wisdom statistical test
5) tracy_widom_statistic = The tracy wisdom test statistic
6) PC = The Principal Component (e.g. PC1 to PC122)

**Dataset S2 (supplementary_indel_pangenes.csv).** Queried GENESPACE pangenomes file for indel region (g17358-g17364).

1) pgID = unique pangenome identifier
2) interpChr = interpolated chromosome in *I. hederacea*
3) interpOrd = interpolated gene-rank order in *I. hederacea*
4) og = orthogroup ID
5) repGene = representative gene
6) genome = genome of the repGene
7) chr = chromosome of the repGene in the species listed in “genome” column
8) start = gene start bp position of the repGene in the species listed in “genome” column
9) end = gene end bp position of the repGene in the species listed in “genome” column
10) Ipomoea_nil = orthologs in *I. nil*
11) Ipomoea_hederacea = orthologs in *I. hederacea*
12) Ipomoea_purpurea = orthologs in *I. purpurea*
13) Ipomoea_triloba = orthologs in *I. triloba*
14) Ipomoea_aquatica = orthologs in *I. aquatica*
15) Ipomoea_cairica = orthologs in *I. carica*

**Dataset S3 (Ihed_all_pheno_GWAS_input20251022.csv).** Phenotype data for homozygote seed families (n=122) measured in common garden experiment. Columns description:

1) Seedfamily = *I. hederacea* seed family
2) Leaf_count1_June15 = First leaf count collected on June 15th 2022
3) Length_1_June16 = First measurement of main vine length collected on June 16th 2022 (cm)
4) Leaf_area = Leaf area ($cm^2$)
5) Leaf_perimeter = Leaf perimeter (cm)
6) Leaf_circularity = Leaf circularity
7) Leaf_AR = Leaf aspect ratio
8) Leaf_roundness = Leaf roundness
9) Leaf_solidity = Leaf solidity
10) leaf_count_2_ June22 = Second leaf count collected June 22nd 2022
11) length_2_June22 = Second measurement of main vine length collected on June 22nd 2022 (cm)
12) Leaf_count_3 = Third leaf count collected collected on June 29th 2022
13) leaf_count_4_July6 = Fourth leaf count collected on July 6th 2022
14) stomatal_conductance = stomatal conductance ($mmol/m^2s$)
15) leaf_temp = leaf temperature (°C)
16) leaf_fresh_weight = leaf fresh weight (g)
17) leaf_turgid_weight = leaf turgid weight (g)
18) leaf_dry_weight = leaf dry weight (g)
19) trichome_count = trichome density
20) relative_water_content = relative water content
21) GWAS_3Lobed_shape = leaf shape as either “heart” or “lobed”

**Dataset S4 (top_hits_arabidopsis.csv).** Top BLAST hits for *I. hederacea* genes found within 250 kb upstream or downstream of the two most significant lobed vs entire GWAS SNPs BLASTed against *A.*

*thaliana* TAIR10 annotation downloaded from NCBI. Columns are labelled similar to BLAST output where *I. hederacea* is the query sequence and *A. thaliana* is the subject. Columns description:

1) salltitles = All subject title(s)
2) stitle = Subject title
3) qseqid = Query Seq-ID which corresponds to *I. hederacea* gene and transcript ID (e.g. g17327.t1)
4) sseqid = Subject Seq-ID
5) pident = Percentage of identical matches
6) length = Alignment length
7) mismatch = Number of mismatches
8) gapopen = Number of gap openings
9) qstart = Start of alignment in query
10) qend = End of alignment in query
11) sstart = Start of alignment in subject
12) send = End of alignment in subject
13) evalue = Expect value
14) bitscore = Bit score

**Dataset S5 (top_hits_ncbi.csv).** Top BLAST hits for *I. hederacea* genes found within 250 kb upstream or downstream of the two most significant lobed vs entire GWAS SNPs BLASTed against the NCBI nr database. Columns are labelled similar to BLAST output. Columns description:

1) salltitles = All subject title(s)
2) stitle = Subject title
3) qseqid = Query Seq-ID which corresponds to *I. hederacea* gene and transcript ID (e.g. g17327.t1)
4) sseqid = Subject Seq-ID
5) pident = Percentage of identical matches
6) length = Alignment length
7) mismatch = Number of mismatches
8) gapopen = Number of gap openings
9) qstart = Start of alignment in query
10) qend = End of alignment in query
11) sstart = Start of alignment in subject
12) send = End of alignment in subject
13) evalue = Expect value
14) bitscore = Bit score

### SI Reference


1. P. Danecek, *et al.*, The variant call format and VCFtools. *Bioinformatics* **27**, 2156–2158 (2011).
2. P. Danecek, *et al.*, Twelve years of SAMtools and BCFtools. *GigaScience* **10**, giab008 (2021).
3. B. E. Campitelli, J. R. Stinchcombe, Natural selection maintains a single-locus leaf shape cline in Ivyleaf morning glory, *Ipomoea hederacea*. *Mol. Ecol.* **22**, 552–564 (2013).
4. H. Cheng, M. Asri, J. Lucas, S. Koren, H. Li, Scalable telomere-to-telomere assembly for diploid and polyploid genomes with double graph. *Nat. Methods* **21**, 967–970 (2024).
5. H. Cheng, G. T. Concepcion, X. Feng, H. Zhang, H. Li, Haplotype-resolved de novo assembly using phased assembly graphs with hifiasm. *Nat. Methods* **18**, 170–175 (2021).
6. H. Cheng, *et al.*, Haplotype-resolved assembly of diploid genomes without parental data. *Nat. Biotechnol.* **40**, 1332–1335 (2022).
7. C. Zhou, S. A. McCarthy, R. Durbin, YaHS: yet another Hi-C scaffolding tool. *Bioinformatics* **39**, btac808 (2023).
8. N. C. Durand, *et al.*, Juicer provides a one-click system for analyzing loop-resolution Hi-C experiments. *Cell Syst.* **3**, 95–98 (2016).
9. N. C. Durand, *et al.*, Juicebox provides a visualization system for Hi-C contact maps with unlimited zoom. *Cell Syst.* **3**, 99–101 (2016).
10. J. M. Flynn, *et al.*, RepeatModeler2 for automated genomic discovery of transposable element families. *Proc. Natl. Acad. Sci.* **117**, 9451–9457 (2020).
11. A. F. A. Smit, R. Hubley, P. Green, RepeatMasker Open-4.0. (2013). Deposited 2015 2013.
12. A. M. Bolger, M. Lohse, B. Usadel, Trimmomatic: a flexible trimmer for Illumina sequence data. *Bioinformatics* **30**, 2114–2120 (2014).
13. H. Dai, Y. Guan, Nubeam-dedup: a fast and RAM-efficient tool to de-duplicate sequencing reads without mapping. *Bioinformatics* **36**, 3254–3256 (2020).
14. Y. Peng, *et al.*, IDBA-tran: a more robust de novo de Bruijn graph assembler for transcriptomes with uneven expression levels. *Bioinforma. Oxf. Engl.* **29**, i326-334 (2013).
15. K. J. Hoff, A. Lomsadze, M. Borodovsky, M. Stanke, "Whole-genome annotation with BRAKER" in *Gene Prediction: Methods and Protocols*, M. Kollmar, Ed. (Springer, 2019), pp. 65–95.
16. F. Tegenfeldt, *et al.*, OrthoDB and BUSCO update: annotation of orthologs with wider sampling of genomes. *Nucleic Acids Res.* **53**, D516–D522 (2025).
17. S. Andrews, FastQC: A quality control tool for high throughput sequence data. (2010). Available at: https://www.bioinformatics.babraham.ac.uk/projects/fastqc/ [Accessed 7 November 2025].
18. P. Ewels, M. Magnusson, S. Lundin, M. Käller, MultiQC: summarize analysis results for multiple tools and samples in a single report. *Bioinformatics* **32**, 3047–3048 (2016).
19. F. Krueger, *et al.*, TrimGalore: v0.6.10. (2023). https://doi.org/10.5281/zenodo.7598955. Deposited 2 February 2023.
20. H. Li, Aligning sequence reads, clone sequences and assembly contigs with BWA-MEM. [Preprint] (2013). Available at: http://arxiv.org/abs/1303.3997 [Accessed 7 November 2025].
21. K. Okonechnikov, A. Conesa, F. García-Alcalde, Qualimap 2: advanced multi-sample quality control for high-throughput sequencing data. *Bioinformatics* **32**, 292–294 (2016).
22. C. C. Chang, *et al.*, Second-generation PLINK: rising to the challenge of larger and richer datasets. *GigaScience* **4**, s13742-015-0047–8 (2015).
23. A. R. Quinlan, I. M. Hall, BEDTools: a flexible suite of utilities for comparing genomic features. *Bioinformatics* **26**, 841–842 (2010).
24. R Core Team, R: A language and environment for statistical computing. (2024). Deposited 2024.
25. Posit Team, Rstudio: Integrated Development Environment for R. (2024). Deposited 2024.
26. H. Wickham, *ggplot2: Elegant graphics for data analysis* (Springer-Verlag New York, 2016).
27. A. Kassambara, ggpubr: "ggplot2" based publication ready plots. (2025). Deposited 2025.
28. T. L. Pedersen, patchwork: The composer of plots. (2025). Deposited 2025.
29. Claus O. Wilke, Brenton M. Wiernik, ggtext: Improved text rendering support for "ggplot2." (2022). Deposited 2022.
30. David Wilkins, ggfittext: Fit text inside a box in "ggplot2." (2024). Deposited 2024.
31. B. Auguie, gridExtra: Miscellaneous functions for "Grid" graphics. (2017). Deposited 2017.
32. B. E. Campitelli, John. R. Stinchcombe, Population dynamics and evolutionary history of the weedy vine *Ipomoea hederacea* in North America. *G3amp58 GenesGenomesGenetics* **4**, 1407–1416 (2014).

33. B. K. Epperson, M. T. Clegg, Spatial-autocorrelation analysis of flower color polymorphisms within substructured populations of Morning Glory (*Ipomoea purpurea*). *Am. Nat.* **128**, 840–858 (1986).
34. D. F. Alvarado-Serrano, M. L. Van Etten, S.-M. Chang, R. S. Baucom, The relative contribution of natural landscapes and human-mediated factors on the connectivity of a noxious invasive weed. *Heredity* **122**, 29–40 (2019).
35. A. J. Stock, B. E. Campitelli, J. R. Stinchcombe, Quantitative genetic variance and multivariate clines in the Ivyleaf morning glory, *Ipomoea hederacea*. *Philos. Trans. R. Soc. B Biol. Sci.* **369**, 20130259 (2014).
36. G. A. Henry, J. R. Stinchcombe, G-matrix stability in clinally diverging populations of an annual weed. *Evolution* **77**, 49–62 (2023).
37. A. K. Simonsen, J. R. Stinchcombe, Quantifying evolutionary genetic constraints in the Ivyleaf Morning Glory, *Ipomoea hederacea*. *Int. J. Plant Sci.* **171**, 972–986 (2010).
38. B. E. Campitelli, J. R. Stinchcombe, Testing potential selective agents acting on leaf shape in *Ipomoea hederacea*: predictions based on an adaptive leaf shape cline. *Ecol. Evol.* **3**, 2409–2423 (2013).
39. B. E. Campitelli, A. J. Gorton, K. L. Ostevik, J. R. Stinchcombe, The effect of leaf shape on the thermoregulation and frost tolerance of an annual vine, *Ipomoea hederacea* (Convolvulaceae). *Am. J. Bot.* **100**, 2175–2182 (2013).
40. Y. K. Singhal, J. A. Boyle, J. R. Stinchcombe, Differences in stomatal conductance between leaf shape genotypes of *Ipomoea hederacea* suggest divergent ecophysiological strategies. *MicroPublication Biol.* (2025). https://doi.org/10.17912/micropub.biology.001528.
41. B. E. Campitelli, A. K. Simonsen, A. Rico Wolf, J. S. Manson, J. R. Stinchcombe, Leaf shape variation and herbivore consumption and performance: a case study with *Ipomoea hederacea* and three generalists. *Arthropod-Plant Interact.* **2**, 9–19 (2008).
42. K. L. Bright-Emlen, “Geographic variation and natural selection on a leaf shape polymorphism in the ivyleaf morning glory (*Ipomoea hederacea*),” Duke University, United States -- North Carolina. (1998).
43. J. A. Boyle, M. E. Frederickson, J. R. Stinchcombe, Genetic architecture of heritable leaf microbes. *Microbiol. Spectr.* **12**, e0061024 (2024).
44. S. Gupta, *et al.*, Interchromosomal linkage disequilibrium and linked fitness cost loci associated with selection for herbicide resistance. *New Phytol.* **238**, 1263–1277 (2023).
